# An ancient viral epidemic involving host coronavirus interacting genes more than 20,000 years ago in East Asia

**DOI:** 10.1101/2020.11.16.385401

**Authors:** Yassine Souilmi, M. Elise Lauterbur, Ray Tobler, Christian D. Huber, Angad S. Johar, David Enard

## Abstract

The current SARS-CoV-2 pandemic has emphasized the vulnerability of human populations to novel viral pressures, despite the vast array of epidemiological and biomedical tools now available. Notably, modern human genomes contain evolutionary information tracing back tens of thousands of years, which may help identify the viruses that have impacted our ancestors – pointing to which viruses have future pandemic potential. Here, we apply evolutionary analyses to human genomic datasets to recover selection events involving tens of human genes that interact with coronaviruses, including SARS-CoV-2, that likely started more than 20,000 years ago. These adaptive events were limited to the population ancestral to East Asian populations. Multiple lines of functional evidence support an ancient viral selective pressure, and East Asia is the geographical origin of several modern coronavirus epidemics. An arms race with an ancient coronavirus, or with a different virus that happened to use similar interactions as coronaviruses with human hosts, may thus have taken place in ancestral East Asian populations. By learning more about our ancient viral foes, our study highlights the promise of evolutionary information to better predict the pandemics of the future. Importantly, adaptation to ancient viral epidemics in specific human populations does not necessarily imply any difference in genetic susceptibility between different human populations, and the current evidence points toward an overwhelming impact of socioeconomic factors in the case of COVID-19.

## Introduction

In the past 20 years, strains of the beta coronavirus genus (family Coronaviridae; Richman et al., 2020) have been behind three major zoonotic outbreaks with grave impacts for human populations (Ou et al., 2020). The first outbreak, commonly known as SARS-CoV (Severe Acute Respiratory Syndrome), originated in China in late 2002 and eventually spread to 30 additional counties where it infected more than 8,000 people and claimed nearly 800 lives (Hoffmann and Kamps, 2003). Four years later, MERS-CoV (Middle East respiratory syndrome coronavirus) affected >2,400 people and caused over 850 deaths, mostly in Saudi Arabia (World Health Organization, 2019). The most recent outbreak began in late 2019 when SARS-CoV-2 – a less virulent but far more contagious strain than those behind the two previous epidemics – emerged in mainland China before spreading rapidly across the rest of the world, triggering an ongoing pandemic (COVID-19) that so far has infected 45 million people and resulted in over one million deaths worldwide (Dong et al., 2020).

The devastation caused by SARS-CoV-2 has inspired a worldwide research effort to develop new vaccines and strategies that aim to curb its impact by determining the factors that underlie its epidemiology. The resulting research has revealed that socioeconomic (e.g. access to healthcare and testing facilities or exposure at work), demographic (e.g. population density and age structure), and personal health factors all play a major role in SARS-CoV-2 epidemiology (Balogun et al., 2020; Sattar Naveed et al., 2020; Scarpone et al., 2020). Additionally, several genetic loci that mediate SARS-CoV-2 susceptibility and severity have been found in contemporary European populations (Ellinghaus et al., 2020; Roberts et al., 2020), one of which contains a genetic variant that increases SARS-CoV-2 susceptibility that likely increased in frequency in the ancestors of modern Europeans after interbreeding with Neanderthals ∼40,000 years ago (Zeberg and Pääbo, 2020). This historical admixture event has led to genetic differences within and between contemporary human populations that directly impact COVID-19 epidemiology – the Neanderthal-derived variant haplotype is now carried by 8% of modern Europeans, but at lower frequencies in African populations whose ancestors did not experience this admixture event – and suggests that evolutionary analyses of human populations may help reveal these genetic differences and ultimately assist in the development of novel drugs and therapies to combat the negative impacts of SARS-CoV-2.

Throughout the evolutionary history of our species, positive natural selection has frequently targeted proteins that physically interact with viruses – e.g. those involved in immunity, or used by viruses to hijack the host cellular machinery (Barreiro et al., 2009; Enard et al., 2016; Sawyer et al., 2005). In the ∼6 million years since the ancestors of humans and chimpanzees separated, selection has led to the fixation of gene variants encoding virus-interacting proteins (VIPs) at three times the rate observed for other classes of genes (Enard et al., 2016; Uricchio et al., 2019). Moreover, strong selection on VIPs has continued in human populations during the past 50,000 years, as evidenced by VIP genes being enriched for adaptive introgressed Neanderthal variants and also selective sweep signals (i.e. selection that drives a beneficial variant to substantial frequencies in a population), particularly around VIPs that interact with RNA viruses, a viral class that includes the coronaviruses (Enard and Petrov, 2018, 2020).

The accumulated evidence suggests that ancient RNA virus epidemics have occurred frequently during the history of our species; however, we currently do not know if selection has made a substantial contribution to the evolution of human genes that interact more specifically with coronaviruses.

Accordingly, here we investigate whether ancient coronavirus epidemics have driven past adaptation within and across modern human populations, by examining if selection signals are enriched within a set of 420 VIPs that interact with coronaviruses (denoted CoV-VIPs; Table S1) across 26 worldwide human populations from the 1000 Genomes Project (1000 Genomes Project Consortium, 2015). These CoV-VIPs comprise 332 SARS-CoV-2 VIPs that were recently identified by high-throughput mass spectrometry (Gordon et al., 2020) and an additional 88 proteins that were manually curated from the coronavirus literature (e.g. SARS-CoV-1, MERS, HCoV-NL63, etc; Table S1; Enard and Petrov, 2018), and form part of a larger set of 5,291 previously published VIPs (SI; Table S1) from multiple viruses known to infect humans (Enard and Petrov, 2018). Our focus upon host adaptation at VIPs is motivated by evidence indicating that these protein interactions are the central mechanism that viruses use to hijack the host cellular machinery, as shown by the strong focus of virologists on these interactions (Enard and Petrov, 2018). Accordingly, VIPs are much more likely to have functional impacts on viruses than proteins not known to interact with viruses (see SI: *Host adaptation is expected at VIPs*). Our enrichment-based approach is expected to be particularly powerful if the ancestors of one or more of the 26 modern human populations were exposed to epidemics driven by coronavirus-like viruses that resulted in selection upon multiple CoV-VIPs (see Discussion). An alternative that we cannot exclude however is that a different type of virus that happens to use similar VIPs as coronaviruses might instead create an enrichment in adaptation signals at CoV-VIPs.

Our analyses of CoV-VIPs find a strong enrichment in sweep signals in these proteins across multiple East Asian populations, which is absent from other human populations. This suggests that an ancient coronavirus epidemic (or another virus using similar VIPs) drove an adaptive response in the ancestors of East Asians, which is in agreement with the current geographic range of the major known animal reservoirs of coronaviruses (Wong et al., 2019). Further, by leveraging ancestral recombination graph approaches (Speidel et al., 2019; Stern et al., 2019) we find that amongst the putatively selected CoV-VIPs, 42 first may have come under selection around 900 generations (∼25,000 years, most likely 20,000 years ago or more) ago and exhibit a coordinated adaptive response that lasted until around 200 generations (∼5,000 years) ago.

By drawing upon other publicly available datasets, we show that the CoV-VIP genes are enriched for anti- and proviral effects and variants that affect COVID-19 etiology in the modern European British population (https://grasp.nhlbi.nih.gov/Covid19GWASResults.aspx). We nevertheless do not investigate in which particular direction, as we cannot expect the British population to be representative of East Asian populations in that respect. We further show that the inferred underlying causal mutations are situated near to regulatory variants active in lungs and other tissues negatively impacted by COVID-19. Taken together, these independent lines of evidence provide support for an ancient coronavirus (or another virus that was using similar interactions) epidemic that emerged more than 20,000 years ago in the ancestors of contemporary East Asian populations, whose genetic signature remains apparent in the genomes of the present-day populations now living in this region.

## Results

### Signatures of adaptation to an ancient epidemic

Viruses have exerted strong selective pressures on the ancestors of modern humans (Enard and Petrov, 2020; Uricchio et al., 2019). Accordingly, we use two population genetic statistical tests that are sensitive to such genetic signatures (i.e. selective sweeps) – nSL (Ferrer-Admetlla et al., 2014) and iHS (Voight et al., 2006) – and which are able to detect genomic regions impacted by strong selection across a wide range of parameters (e.g. different starting and end frequencies of the selected allele). Both statistics also have the advantage of being insensitive to background selection (Enard et al., 2014; Schrider, 2020), thereby reducing the potential impact of false positives in our analyses.

After scanning each of the 26 populations for signals of selection, we apply an enrichment test that was previously used to detect enriched selection signals in RNA VIPs in human populations (Enard and Petrov, 2020). Briefly, for each population and selection statistic, we rank all genes based on the average selection statistic score observed in genomic windows ranging from 50kb to 2Mb (Methods). Different windows sizes are used because smaller windows tend to be more sensitive to weaker sweeps, whereas larger windows tend to be more sensitive to stronger sweeps (Enard and Petrov, 2020; Methods). After ranking the gene scores, we estimate an enrichment curve (Figure 1) for gene sets ranging from the top 10 to 10,000 ranked loci (Methods). The significance of the whole enrichment curve is then calculated using a genome block-randomization approach that accounts for the genomic clustering of neighboring CoV-VIPs, and provides an unbiased false positive risk for the whole enrichment curve (FPR) by re-running the entire enrichment analysis pipeline on block-randomized genomes (Enard and Petrov, 2020; Methods). For our control gene set, we use protein-coding genes situated at least 500kb from CoV-VIPs to avoid overlapping the same sweep signals. Additionally, genes in the control sets are chosen to have similar characteristics as the CoV-VIPs (e.g. similar recombination rates, density of coding and regulatory sequences, percentage of immune genes, percentage of genes that interact with bacteria; see Methods for the complete list of factors) to ensure that any detected enrichment is virus-specific rather than due to a confounding factor (Enard and Petrov, 2020). Choosing controls far away and that match multiple potential confounding factors has the effect of shrinking the pool of potential control genes, which can affect the variance and also the representativity of this pool as a null control. The possible impacts of the size of the control pool are however fully taken into account in the FPR estimated with block-randomized genomes (Enard and Petrov, 2020; Methods). Finally, we also exclude the possibility that functions other than viral interactions might explain our results by running a Gene Ontology analysis (Gene Ontology Consortium, 2015; SI; Tables S2, S3 and Figure S1).

**Figure 1.**
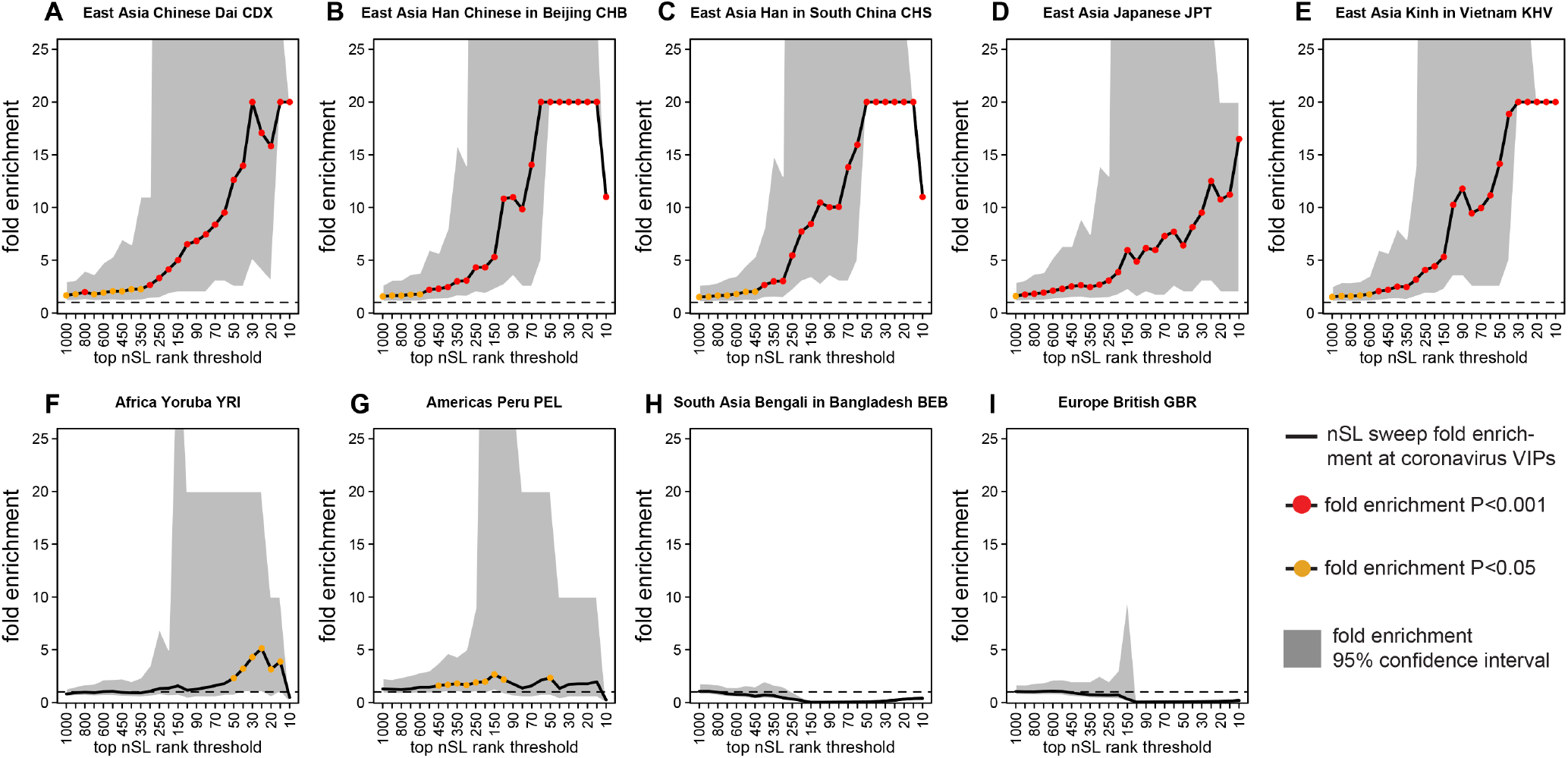
Coronavirus VIPs nSL ranks enrichment. A,B,C,D,E are East Asian populations, F,G,H,I are populations from other continents. The y axis represents the bootstrap test (Methods) relative fold enrichment of the number of genes in putative sweeps at CoV-VIPs, divided by the number of genes in putative sweeps at control genes matched for multiple confounding factors. The x axis represents the top rank threshold to designate putative sweeps. Black full line: average fold enrichment over 5,000 bootstrap test control sets. Fold enrichments greater than 20 are represented at 20. Grey area: 95% confidence interval of the fold enrichment over 5,000 bootstrap test control sets. The rank thresholds where the confidence interval lower or higher fold enrichment has a denominator of zero are not represented (For example, graph B, top 10 rank threshold). Lower confidence interval fold enrichments higher than 20 are represented at 20 (for example, graph B, top 30 rank threshold). Red dots: bootstrap test fold enrichment P<0.001. Orange dots: bootstrap test fold enrichment P<0.05. Note that the bootstrap test p-values are not the same as the whole curve enrichment false positive risk (FPR) estimated using block-randomized genomes on top of the bootstrap test (Methods).

Applying this approach to each of the 26 human populations from the 1,000 genomes dataset, we find a very strong enrichment of sweep signals in CoV-VIPs across all top-ranked gene set sizes that is specific to the five East Asian populations (whole enrichment curve for nSL and iHS combined FPR=2.10^−4^; Figures 1 & S2; Methods). No enrichment is observed for populations from other continental regions, including in neighboring South Asia (whole enrichment curve for nSL and iHS combined FPR>0.05 in all cases; Figures 1 & S2). Further, no enrichment is detected for VIP sets for 17 other viruses in East Asian populations (whole enrichment curve for nSL and iHS separately or combined, P>0.05 in all cases; Figures S3 & S4). Taken together, these results suggest that coronaviruses, or another type of viruses that used similar interactions with human hosts, have driven ancient epidemics in ancient human populations that are ancestral to modern East Asians. This enrichment is unlikely to have been caused by any other virus represented in our set of 5,291 VIPs, but we still cannot exclude that a currently unknown type of virus that happened to use similar VIPs as coronaviruses could have been involved instead (Table S1). The enrichment is most substantial for the top-ranked gene sets ranging between the top 10 and top 1,000 loci (Figure 1; whole enrichment curve FPR=3.10^−6^ for nSL, FPR=4.10^−3^ for iHS, FPR=6.10^−5^ for iHS and nSL combined), and is particularly strong for the top 200 loci in large windows (1 Mb) where a four-fold enrichment is observed for both nSL and iHS statistics (pertaining to between 10 to 13 selected CoV-VIPs amongst the top 200 ranked genes; Table S4). This suggests that strong selection targeted multiple CoV-VIPs in the common ancestors of modern East Asian populations. That the selected haplotype structures are detected by both the iHS and nSL methods suggests that they are unlikely to have occurred prior to 30,000 years ago, as both nSL and iHS have little power to detect adaptive events arising before this time point in human evolution (Sabeti et al., 2006)

### An ancient epidemic in the ancestors of East Asians starting more than 20,000 years ago

To further test the existence of an ancient viral epidemic in the ancestors of East Asians, we use a recent ancestral recombination graph (ARG)-based method, Relate (Speidel et al., 2019), to infer the timing and trajectories of selected loci for the CoV-VIPs. If the selective pressure responsible for the multiple independent selection events at CoV-VIPs was relatively sudden as expect from a new epidemic, then these selection events should have started independently around the same time. By estimating ARGs at variants distributed across the entire genome, Relate can reconstruct coalescent events across time and detect genomic regions impacted by positive selection, while explicitly controlling for historical variation in population demography. To approximate the start time of selection, Relate estimates the first historical time point that a putatively selected variant had an observable frequency unlikely to be equal to zero (Methods). We use this approximation as the likely starting time of selection, although we note that this method does not account for selection on standing variants that had non-zero frequencies at the onset of selection (Methods). Additionally, we use the iSAFE software – which enables the localization of selected mutations (Akbari et al., 2018) – along with a curated set of regulatory variants (expression QTLs; eQTLs) from the eGTEx Project (2017) to help identify the likely causal mutations in the selected CoV-VIP genes. There is good evidence that the majority of adaptive mutations in the human genome are regulatory mutations (Enard et al., 2014; Kudaravalli et al., 2009; Nédélec et al., 2016; Quach et al., 2016) and, accordingly, we find that iSAFE peaks are significantly closer to GTEx eQTLs proximal to CoV-VIP genes than expected by chance (iSAFE peak proximity test, P<10^−9^; Methods). Therefore, for each CoV-VIP gene, we choose a variant with the lowest Relate p-value (<10^−3^; Methods) that is situated at or close to a GTEx eQTL associated with the focal gene to estimate the likely starting time of selection for that gene (Methods; Figure S5).

Using this approach, we observe 42 CoV-VIPs (Table S5 and Figure S5) with selection starting times clustered around a peak 870 generations ago (∼200 generations wide, potentially due to noise in our estimates; Figure 2). While this amounts to about four times more selected CoV-VIP genes than were detected using either nSL or iHS (both detected around ten CoV-VIPs amongst the top 200 ranked genes; Table S4) this is not unexpected as Relate has more power to detect selection events than nSL and iHS when the beneficial allele is at intermediate frequencies at the point of measurement (typically <60%; Figure 3; Enard and Petrov, 2020; Ferrer-Admetlla et al., 2014; Voight et al., 2006). The relatively tight temporal clustering of starting times forms a highly significant peak (peak significance test P=2.3.10^−4^; Figure 2) when comparing the observed clustering of CoV-VIPs start times with the distribution of inferred start times for randomly sampled sets of genes (Methods). Note that this peak significance test is gene clustering-aware (Methods). Further, this significance test is not biased by the fact that CoV-VIPs are enriched for sweep signals, as the test remains highly significant (P=1.10^−4^) when using random control sets with comparable high-scoring nSL statistics (Methods). This suggests that the tight temporal clustering of selection events is a specific feature of the CoV-VIPs, rather than a confounding aspect of any gene set similarly enriched for sweeps.

**Figure 2.**
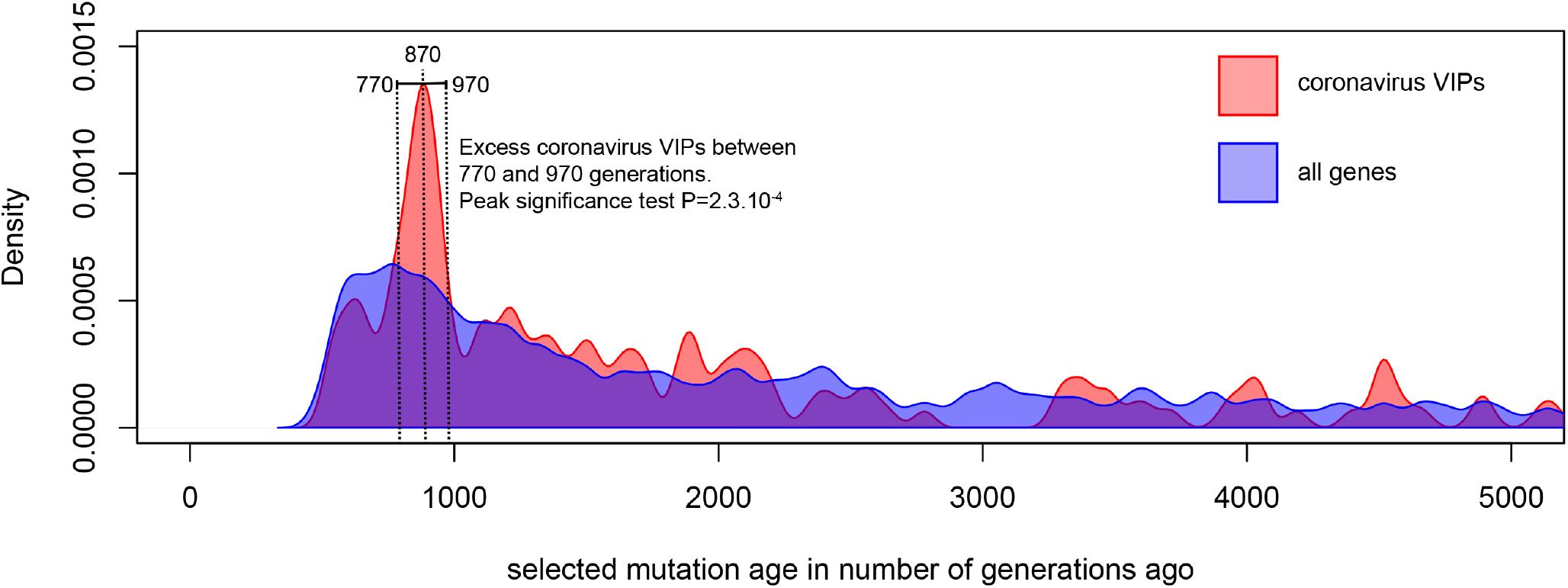
Timing of selection at CoV-VIPs. The figure shows the distribution of selection start times at CoV-VIPs (pink distribution) compared to the distribution of selection start times at all loci in the genome (blue distribution). Details on how the two distributions are compared by the peak significance test, and how the selection start times are estimated with Relate, are provided in Methods.

**Figure 3.**
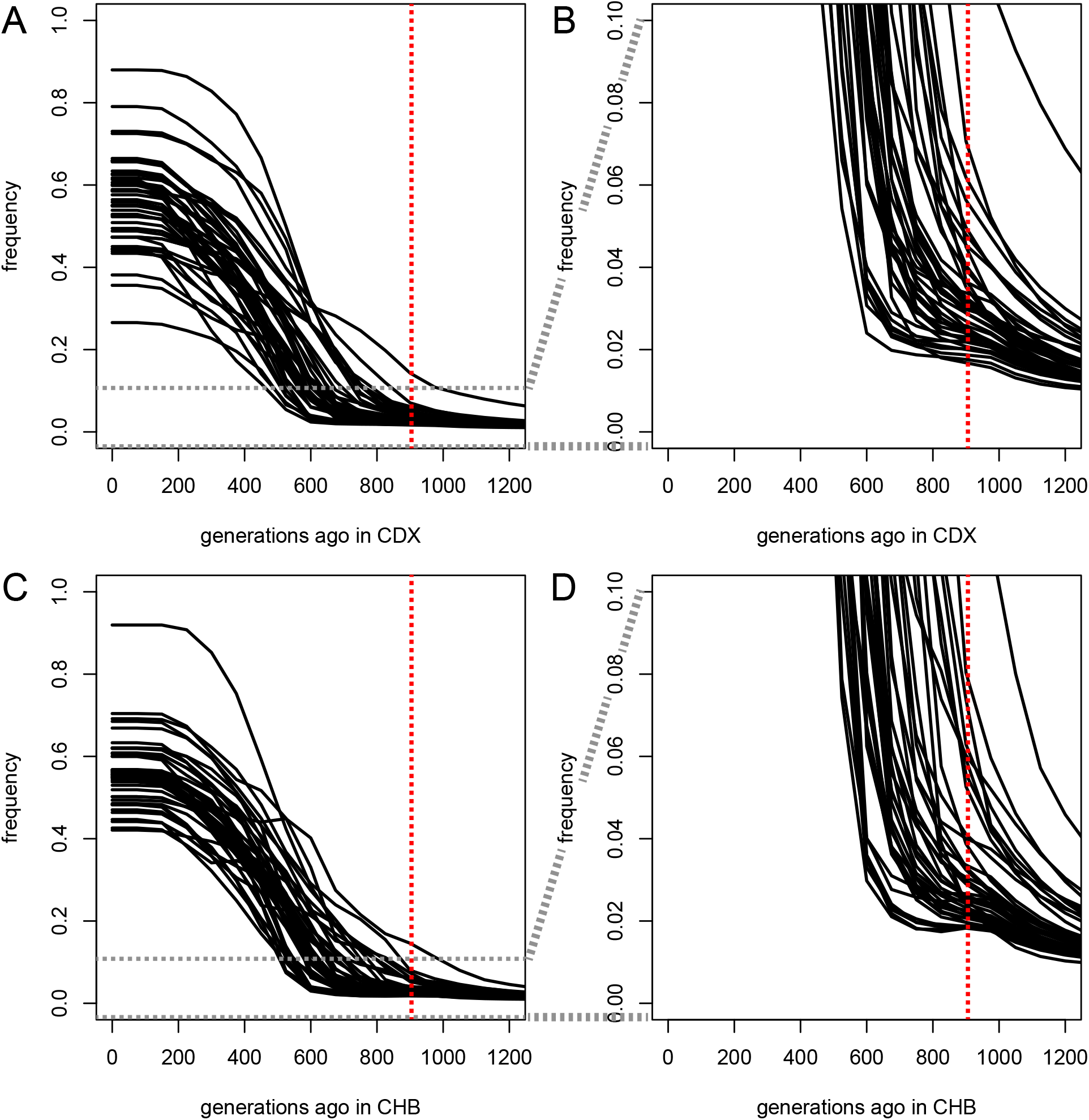
Selected CoV-VIPs allele frequency trajectories over time estimated by CLUES in East Asia. Each frequency trajectory is for one of the 42 Relate selected mutations at CoV-VIPs within the peak around 900 generations ago (Methods). A) Frequency trajectories in the Chinese Dai CDX 1,000 Genomes population. B) Same, but zoomed-in from frequencies 0 to 10%. C) Frequency trajectories in the Han Chinese from Beijing CHB 1,000 Genomes population. D) Same, but zoomed-in from frequencies 0 to 10%.

The genes with clustered selection starting times around 900 generations ago are enriched in strong nSL signals, as shown by running the peak significance test using only CoV-VIPs and controls with strong nSL signals (Figure S6). Conversely, the peak disappears when restricting this test to weaker nSL signals (P=0.53 when using the lowest 50% of nSL statistics; Methods). Importantly, our estimates of the timing of selection are not biased by our use of methods that rely on selected variants not being fixed in the population at the time of genome sampling (i.e. Relate). When rerunning our analytical pipeline focusing only on strong candidate loci according to Tajima’s D (Tajima, 1989), a statistic developed to detect recently completed sweeps (i.e. fixed mutations), we observe the same clustering of selection events starting around 900 generations ago (Figure S7). Further, the remaining 382 CoV-VIPs that are not part of this temporal cluster around 900 generations ago are not more likely to have significant Tajima’s D values than controls (whole enrichment curve P=0.07). Consequently, our results are consistent with the emergence of a viral epidemic ∼900 generations, or ∼25,000 years (900 generations * 28 years per generation; Moorjani et al., 2016), ago that drove a burst of strong positive selection in the ancestors of East Asians, which may represent a genetic record of a multi-generational viral epidemic amongst the 26 human populations tested here.

Although selective pressures other than a coronavirus or another unknown type of virus with similar host interactions might also contribute to these patterns, we note that the signal is restricted specifically at CoV-VIPs and none of 17 other viruses that we tested exhibit the same temporal clustering ∼900 generations ago in East Asia (peak significance test P>0.05 in all cases; Methods). Further, this test remained highly significant when retesting the temporal clustering of CoV-VIPs using only other RNA VIPs as the control set (P=4.10^−4^; Table S1), consistent with the clustered selection signals being a coordinated adaptive response to a coronavirus or another virus using similar host interactions.

### Strong selection drove coordinated changes in multiple CoV-VIP genes over 20,000 years

To learn more about the likely start and duration of the selection pressure acting on the ancestors of East Asians, we use CLUES (Stern et al., 2019) to infer allele frequency trajectories and selection coefficients for the inferred beneficial mutations proximal to the 42 CoV-VIP genes with selection starting 900 generations ago according to Relate (Figure 3). CLUES uses the temporal variation in population size and coalescence rates inferred by Relate to reconstruct frequency trajectories while taking demographic fluctuations into account. Our observation of sweep signals at 42 CoV-VIP genes in the ancestors of East Asians suggests that the putative underlying viral epidemic likely spanned many generations (i.e. the time needed for selection to drive initially rare alleles to intermediate/high frequencies). Accordingly, we anticipate that selection was probably strongest when the naive host population was first infected by the virus, before gradually waning as the host population adapted to the viral pressure (Hayward and Sella, 2019). Similarly, a decrease in the virulence of the virus over time, a phenomenon that has been reported during the long term bouts of host-virus coevolution (Best and Kerr, 2000), would also result in the gradual decrement of selection coefficients across time. Hence, for each of the 42 CoV-VIPs predicted to have started coming under selection ∼900 generations ago, we use CLUES to estimate the selection coefficient in two successive time-intervals (between 1,000 and 500 generations ago, and from 500 generations ago to the present), predicting that selection would be stronger in the oldest interval. We note that a 500 generations interval was reported as the approximate timespan that CLUES provides reliable estimates for humans (Stern et al., 2019); using smaller generations intervals, we would run the risk of getting overly noisy selection coefficient estimates based on too few coalescent events. However, 500 generations intervals are not adequate to obtain reasonable estimates of the precise duration of the selective pressure (Stern et al., 2019), so we do not attempt to estimate this parameter here, and we simply try to compare the two time periods with each other. Also, because CLUES uses a computationally intensive algorithm when following the recommendations of Stern et al. (2020), we base our estimates on only two of the five East Asian populations (i.e. Dai and Beijing Han Chinese; Figure 3A, B and 3C, D, respectively).

CLUES infers frequency trajectories that are more complex than a simple, clear, abrupt jump in frequency 900 generations ago. Instead, the estimated frequency trajectories (Figure 3A,B,C,D) suggest that 900 generations ago is the approximate time when the bulk of the selected variants reached a frequency of a few percent or more, and approximately when there is an acceleration in the frequency increase (Figure 3B, D). This might correspond to the transition between the establishment and exponential phases of the sweeps, and might imply that the selective pressure is older than 900 generations. The initially flatter, slower increases in frequency, lasting sometimes up to 600 generations ago for some variants, are compatible with either co-dominant or recessive alleles, and likely exclude dominant alleles that would start increasing in frequency more abruptly. Interestingly, this would be in good agreement with the rarity of dominant eQTLs in GTEx, if selected variants were indeed regulatory (GTEx Project, 2017). Although the flat, slow starts of frequency increases make it hard to pinpoint when selection started exactly, the vast majority of the selected alleles appear to have reached 5% or higher frequencies by 600 generations, thus making it highly unlikely that the selective pressure would have started 600 or less generations ago. Frequency trajectories estimated in the Yoruba African population (Figure 4A) or the British European population (Figure 4B) also show very low frequencies 900 generations ago. The selected variants in East Asia are found nowadays at very low frequencies especially in Africa (Table S6). This implies that they are substantially older than when selection started in East Asia, which may then be described as selection on low frequency standing variation. Intriguingly, some variants rise in frequency (up to 40% frequency at most) in Europe mostly after 800 generations ago. A small number of variants in Africa increase in frequency (up to 30% frequency at most) after 600 generations ago.

**Figure 4.**
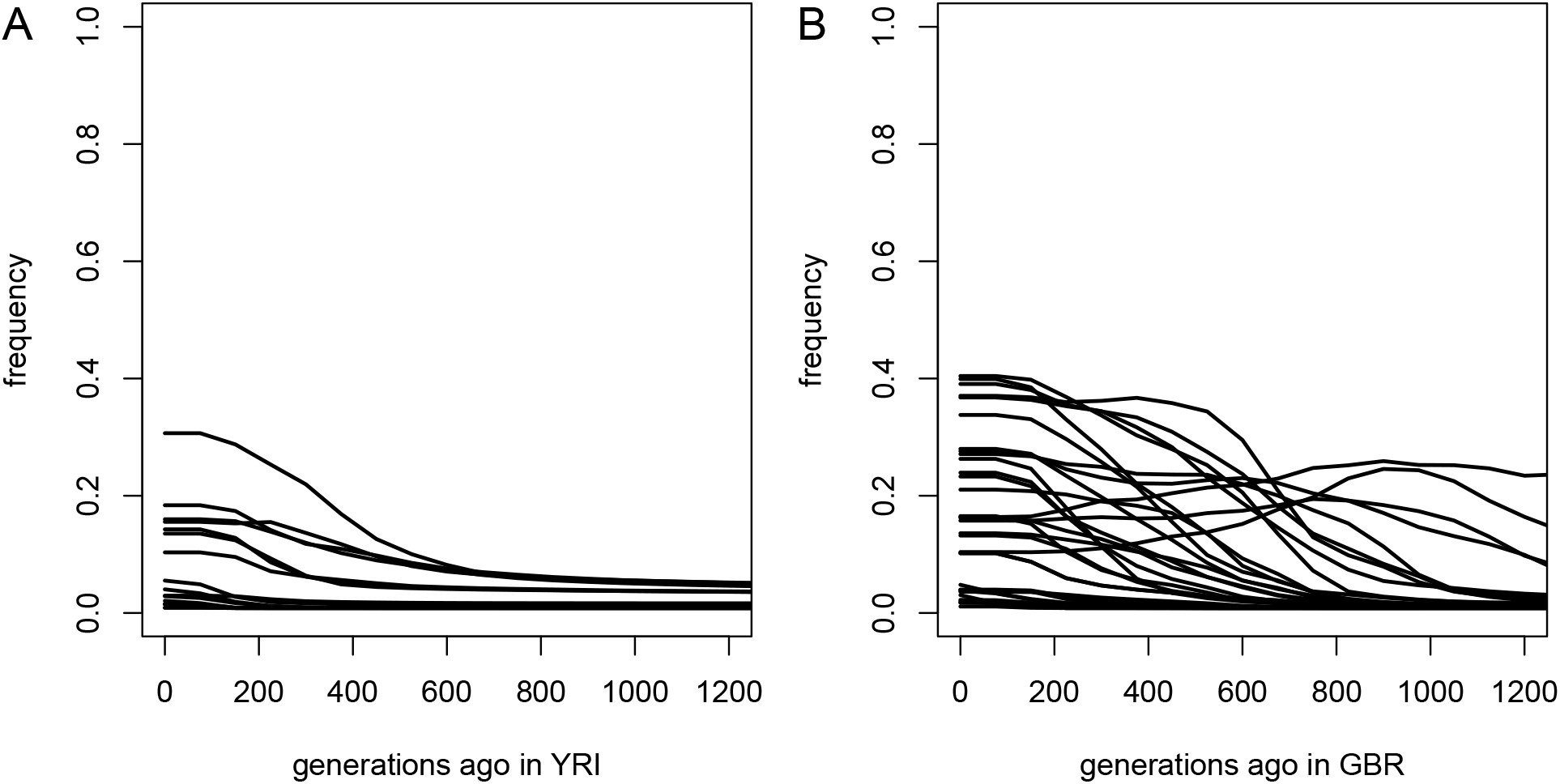
Selected CoV-VIPs allele frequency trajectories over time estimated by CLUES in Africa (Yoruba) and Europe (British) Same as Figure 3. A) Yoruba population. The graph includes 17 frequency trajectories, the 25 other alleles selected in East Asia being absent in the Yoruba sample (but not Africa overall, see Table Sx) B) British population. The graph includes 35 frequency trajectories, the other seven alleles selected in East Asia being absent in the British sample.

The selected mutations are estimated to have continually increased in frequency in East Asia until ∼200 generations (approximately 5,000 years) ago, after which they remained relatively stable (Figure 3A, C). Accordingly, CLUES estimates very high selection coefficients in the interval between 1,000 and 500 generations ago (Dai average *s* = 0.034, Beijing Han average s = 0.042; Figure 5A, B), but much weaker selection coefficients from 500 generations ago up to the present (Dai average *s* = 0.002, Beijing Han average *s* = 0.003; Figure 5A, B). These patterns are consistent with the appearance of a strong selective pressure that triggered a coordinated adaptive response across multiple independent loci, which waned through time as the host population adapted to the viral pressure and/or as the virus became less virulent.

**Figure 5.**
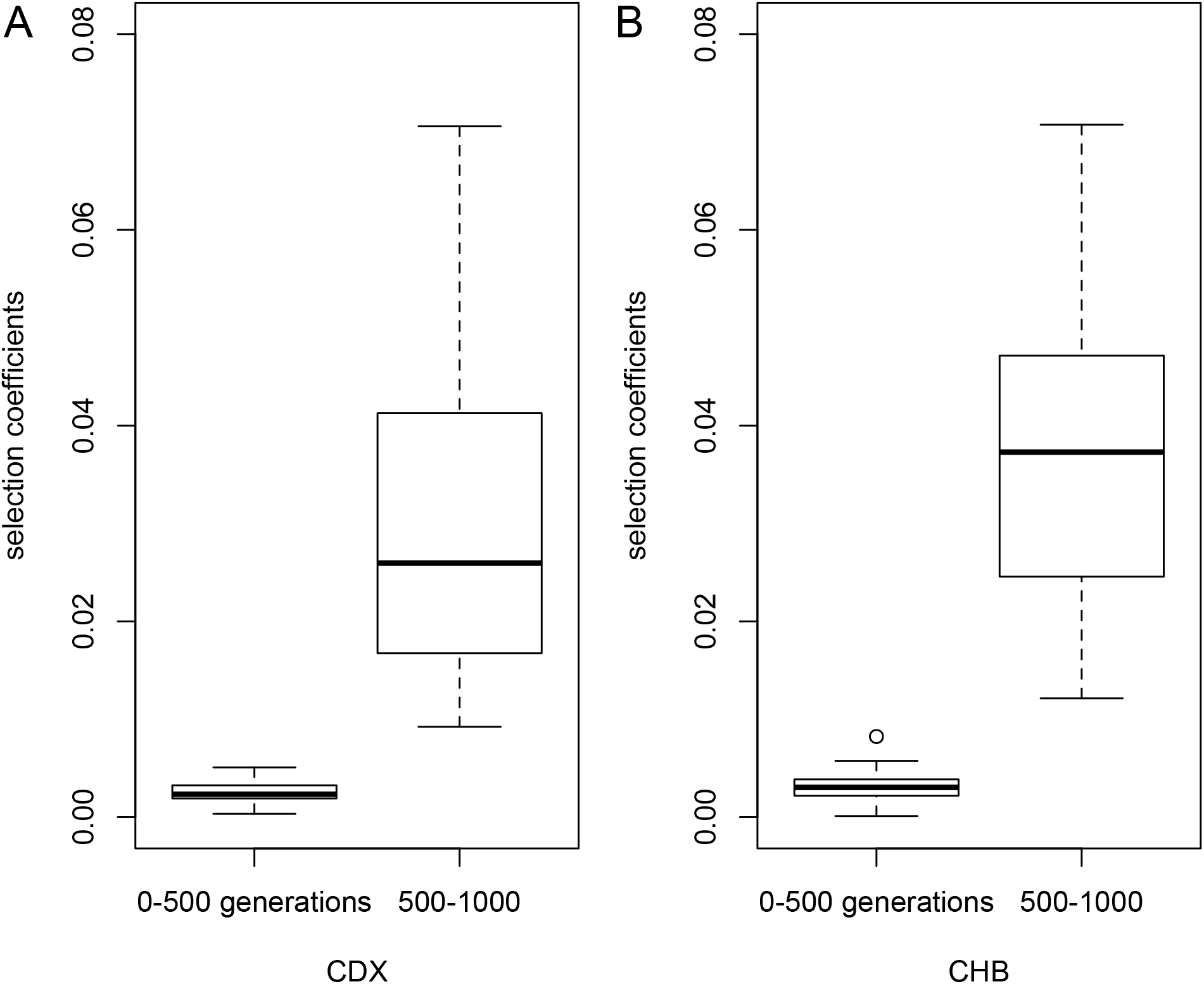
Coronavirus selected VIPs selection coefficients estimated by CLUES. This figure shows classic R boxplots of selected coefficients at the 42 Relate selected mutations within the peak around 900 generations ago (Methods). A) Selection coefficients in the Chinese Dai CDX 1,000 Genomes population. B) Selection coefficients in the Han Chinese from Beijing CHB 1,000 Genomes population. Left: average selection coefficients between 0 and 500 generations ago. Right: average selection coefficients between 500 and 1,000 generations ago.

### Selected CoV-VIPs are enriched for antiviral and proviral factors

To further clarify that an ancient viral epidemic caused the strong burst of selection we observe in the ancestors of East Asians, and not another ecological pressure acting on the same set of genes, we test if the 42 selected CoV-VIPs are enriched for genes with antiviral or proviral effects relative to other CoV-VIPs (i.e. loci that are known to have a detrimental or beneficial effect on the virus, respectively). Because the relevant literature for coronaviruses is currently limited – which also applies to the relatively recent SARS-CoV-2 virus – we extend our set of anti- and proviral loci beyond those associated with coronaviruses to include loci reported for diverse viruses with high confidence from the general virology literature (see SI: *Host adaptation is expected at VIPs*; Table S1). We find that 21 (50%) of the 42 CoV-VIPs that came under selection ∼900 generations ago have high-confidence anti- or proviral effects (vs. 29% for all 420 CoV-VIPs), a significant inflation in anti- and proviral effects (hypergeometric test P=6.10^−4^) that further supports our claim that the underlying selective pressure was most likely a viral epidemic. This overlap of antiviral and proviral effects between different viruses also implies that an unknown virus that happened to use similar VIPs as coronaviruses could have indeed been responsible.

### Selected mutations lie near regulatory variants active in SARS-CoV-2 affected tissues

Coronavirus infections in humans are known to have pathological consequences for specific bodily tissues, whereby we investigate if the genes targeted by selection in the ancestors of East Asians are also enriched for regulatory functions in similar tissues. In light of our finding that many putative causal mutations in CoV-VIPs were proximal to eQTLs, we investigate whether selected mutations are situated closer to eQTLs for a given tissue than expected by chance, as this would indicate that the tissue was negatively impacted by the virus (prompting the adaptive response). Note that the GTEx eQTLs we use are not specific to a single tissue (eQTLs are rarely so in general), and are shared between tissues. However, each tissue still has its own specific combination of eQTLs, thus making the results at each tissue not completely redundant. Briefly, we estimate a proximity-based metric that quantifies the distance between the location of the causal mutation estimated by iSAFE and the tissue-specific eQTLs for the 42 loci that likely started coming under selection ∼900 generations ago, and compare this to the same distances observed amongst randomly sampled sets of CoV-VIPs (Figure 6; Methods).

**Figure 6.**
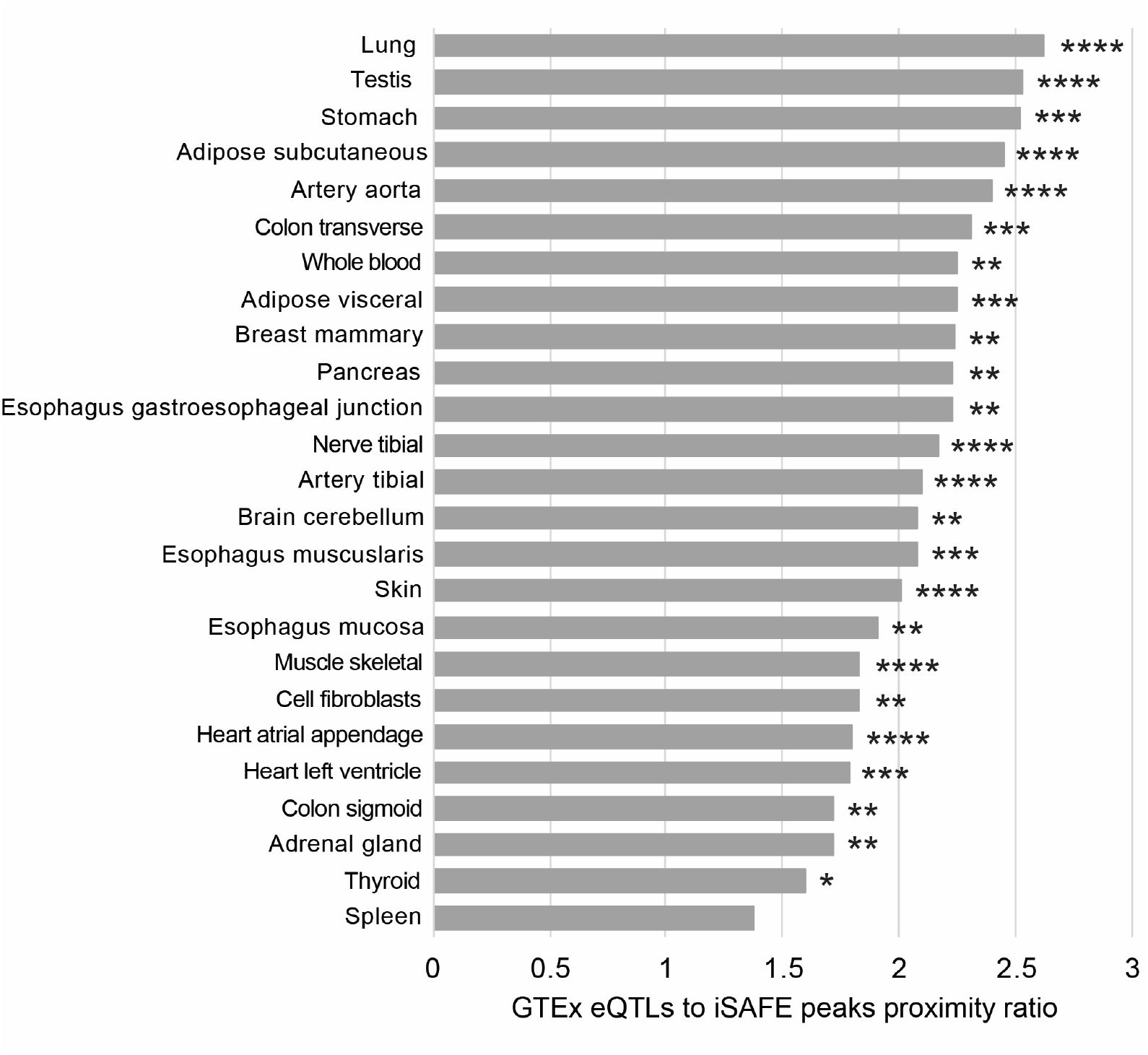
Proximity of selection signals to GTEx eQTLs at the 42 selected CoV-VIPs compared to random CoV-VIPs. The histogram shows how close selection signals localized by iSAFE peaks are to the GTEx eQTLs from 25 different tissues, at peak-VIPs compared to randomly chosen CoV-VIPs (Methods). How close iSAFE peaks are to GTEx eQTLs compared to random CoV-VIPs is estimated through a proximity ratio. The proximity ratio is described in the Methods. It quantifies how much closer iSAFE peaks are to eQTLs of a specific GTEx tissue, compared to random expectations that take the number and structure of iSAFE peaks, as well as the number and structure of GTEx eQTLs into account (Methods). Four stars: proximity ratio test P<0.0001. Three stars: proximity ratio test P<0.001. Two stars: P<0.01. One star: P<0.05. Note that lower proximity ratios can be associated with smaller p-values for tissues with more eQTLs (due to decreased null variance; for example, skeletal muscle vs. pancreas).

Using this approach, we find that GTEx lung eQTLs lie closer to predicted causal mutations amongst the 42 putative selected loci than for any other tissue (P=3.10^−5^; Figure 6). Several additional tissues known to be negatively affected by coronavirus – blood and arteries (Bao et al., 2020; Grosse et al., 2020), adipose tissue (Michalakis and Ilias, 2020) and the digestive tract (Elmunzer et al., 2020) – also exhibit closer proximities between putative causal loci and tissue-specific eQTLs than expected by chance (Figure 6). Interestingly, the spleen shows no tendency for eQTLs to lie closer to selected loci than expected around 900 generations ago compared to other evolutionary times, perhaps because the spleen is replete with multiple types of immune cells that might be more prone to more regular adaptation in response to diverse pathogens over time, and less prone to adaptive bursts restricted over time in response to a specific pathogen (Quintana-Murci, 2019). Note that tissues with more eQTLs tend to have more significant p-values. For example skeletal muscle has a lower proximity ratio than stomach but also a lower p-value due to the higher statistical power provided by more eQTLs. Our results indicate that the tissues impacted in the inferred viral epidemic in ancestors of East Asians match those pathologically affected by the SARS-CoV-2 infection in contemporary populations, providing further evidence that this ancient infection might have been a coronavirus or another type of virus that used similar host interactions.

### Coronavirus VIPs are enriched for SARS-CoV-2 susceptibility and COVID-19 severity loci

Our results indicate that many of the selected CoV-VIPs now sit at intermediate to high frequencies in modern East Asian populations. Accordingly, we anticipate that these segregating loci should make a measurable contribution to the inter-individual variation in SARS-CoV-2 susceptibility and (COVID-19) severity amongst contemporary populations in East Asia, and predict that such loci would be readily detectable in a reasonably-powered genome wide association study (GWAS) investigating these traits in East Asian populations. While such a scan has yet to be reported for a large East Asian cohort, two GWASs were recently released that used sizable British cohorts to investigate SARS-CoV-2 susceptibility (1,454 cases and 7,032 controls; henceforth called the susceptibility GWAS) and severity (325 cases [deaths] versus 1,129 positive controls; henceforth called the severity GWAS) (data from the UK Biobank; Sudlow et al., 2015; https://grasp.nhlbi.nih.gov/Covid19GWASResults.aspx). Because we use a different population than the ones where we found selection, we only ask, as a form of functional validation of a viral pressure, if there is an overlap between the selected loci in East Asia and stronger COVID-19 GWAS hits in the UK Biobank cohort. We do not look at all at the directionality or the size of effects, as it is dubious that those would be transposable between populations. This also means that we make no claim at all here about any decrease or increase of virus susceptibility in any given human population compared to others. Furthermore, we use the UK-Biobank cohort instead of the complete COVID-19 Host Genetics Initiative meta-GWAS data (https://www.covid19hg.org/; The COVID-19 Host Genetics Initiative, 2020), to avoid population stratification to the best extent possible (a legitimate concern with a trait clearly affected by environmental factors).

While we are unable to precisely identify the causal variants for the selected CoV-VIP genes observed in the ancestors of East Asians – nor would these variants necessarily occur as outliers in a GWAS conducted on the British population – we note that it is possible that other variants in the same CoV-VIP genes may also produce variation in SARS-CoV-2 susceptibility and severity amongst modern British individuals.

By contrasting variants in CoV-VIPs against those in random sets of genes, we find that variants in CoV-VIPs have significantly lower p-values for both the susceptibility GWAS and severity GWAS than expected (simple permutation test P<10^−9^ for both GWAS tests; Methods). More importantly, the 42 CoV-VIPs from the selection event starting ∼900 generations ago have even lower GWAS p-values compared to other CoV-VIPs (P=0.0015 for susceptibility GWAS and P=0.023 for severity GWAS; Methods). This result indicates that the selected genes inferred in our study might contribute to individual variation in COVID-19 etiology in modern human populations in the UK, providing further evidence that a coronavirus or another virus with similar host interactions may have been the selection pressure behind the adaptive response we observe in the ancestors of East Asians. Notably, the strongest GWAS hits identified by the COVID-19 Host Genetics Initiative (listed at https://www.covid19hg.org/publications/) do not overlap with the 42 CoV-VIPs selected in East Asia. We note however that we do not necessarily expect the strongest GWAS hits in Europe to be strong hits in other populations. In addition, although adaptation implies a functional genetic effect, a genetic effect does not necessarily mean it has adaptive potential. The lack of overlap with the strongest COVID-19 Host Genetics Initiative hits is therefore not necessarily very surprising. It also does not take away the fact that we found an enrichment in stronger GWAS hits on average at CoV-VIPs and especially at selected CoV-VIPs.

### Selected CoV-VIP genes include multiple known drug targets

Our analyses suggest that the 42 CoV-VIPs identified as putative targets of an ancient coronavirus (or another virus using similar host interactions) epidemic might play a functional role in SARS-CoV-2 etiology in modern human populations. We find that four of these genes (*SMAD3, IMPDH2, PPIB, GPX1*) are targets of eleven drugs being currently used or investigated in clinical trials to mitigate COVID-19 symptoms (Methods). While this number is not higher than expected when compared to other CoV-VIPs (hypergeometric test P>0.05), we note that most of the 42 genes identified here have yet to be the focus of clinical trials for SARS-CoV-2-related drugs. In addition to the four selected CoV-VIP genes targeted by coronavirus-specific drugs, five additional selected CoV-VIPs are targeted by multiple drugs to treat a variety of non-coronavirus pathologies (Table S7). This raises the possibility that such drugs could be repurposed for therapeutic use in the current SARS-CoV-2 pandemic. Indeed, an additional six of the 42 selected CoV-VIPs have been identified by (Finan et al., 2017) as part of the “druggable genome” (Table S7).

## Discussion

By scanning 26 diverse human populations from five continental regions for evidence of strong selection acting on genes that interact with coronavirus strains (CoV-VIPs), we identified a set of 42 CoV-VIPs exhibiting a coordinated adaptive response that likely emerged more than 20,000 years ago (Figure 2). This pattern was unique to the ancestors of East Asian populations (as classified by the 1,000 Genomes, including South East Asia with the Kinh in Vietnam), being absent from any of the 21 non East-Asian human populations tested here. By using ARG methods to reconstruct the trajectories of selected alleles, we show that this selection pressure produced a strong response across the 42 CoV-VIP genes that gradually waned and resulted in the selected loci plateauing at intermediate frequencies. Further, we demonstrate that this adaptive response is likely the outcome of a multigenerational viral epidemic, as attested by the clustering of putatively selected loci around variants that regulate tissues known to exhibit COVID-19-related pathologies, and the enrichment of variants associated with SARS-CoV-2 susceptibility and severity, as well as anti- and proviral functions, amongst the 42 CoV-VIP genes selected starting around 900 generations ago.

An important limitation of our study is that some of our analyses rely upon comparative datasets that were generated in contemporary human populations that have different ancestries than the East Asian populations where the selected CoV-VIP genes were detected. In particular, both of the eQTL and GWAS datasets come from large studies that are primarily focused on contemporary populations from Europe, and none of the five European populations in our study exhibit the selection signals observed in the genomes of East Asians. Accordingly, more direct confirmation of the causal role of 42 CoV-VIP genes in COVID-19 etiology will require the appropriate GWAS to be conducted in East Asian populations. The detection of genetic associations amongst the 42 CoV-VIPs in a GWAS on contemporary East Asians would provide further evidence that one or more coronaviruses, or another virus using similar interactions, comprised the selection pressure that drove the observed adaptive response. Moreover, a high-powered GWAS in East Asian populations would be required to identify the loci that currently impact individual variation in COVID-19 etiology in East Asian individuals. Because of these limitations, and because it would be extremely difficult to control for all the other factors that differ across the world (including socioeconomic factors), our results do not represent evidence for any difference in either increased or decreased genetic susceptibility in any human population.

### Insights into ancient viral epidemics from modern human genomes

A particularly salient feature of the adaptive response observed for the 42 CoV-VIPs is that selection appears to be acting continuously over a ∼20,000 years period, with the caveat that the start of selection is complex to pinpoint as shown by the analysis of the selected alleles frequency trajectories (Figure 3). The activity of a viral pressure over such an extensive time period is not consistent with epidemics that started in recorded human history, which tend to be circumscribed to a few generations. A possible hypothesis is that the viral pressure remained present throughout the 20,000 year period, but was only initially strong enough to qualify as a full-blown pandemic in the commonly understood sense, before becoming less severe over time as a consequence of host adaptation and/or a reduction in virulence. As this manuscript was in the final stages of preparation, the first host-virus interactomes were published for SARS-CoV-1 and MERS-CoV, which exhibit an extensive overlap with the SARS-CoV-2 interactome used in the present study (Gordon et al., 2020). This suggests that coronaviruses share a broad set of host proteins that they interact with, which should also apply to ancient coronaviruses. These patterns are consistent with one or more coronaviruses driving selection events in East Asian prehistory that produced the signals that we report here. That said, and as already mentioned, we cannot exclude that another, currently unknown type of viruses might have been responsible, that used the same interactions as coronaviruses with human proteins. The cumulated evidence in this study still clearly points towards an ancient viral selective pressure.

Further validation of the historical trajectories of the causal mutations at selected genes is still needed, including more finely resolved temporal and geographic patterns that could be derived from ancient DNA sampled from across East Asia that span the human occupation of this region; however, the requisite ancient samples are lacking at the moment. Nonetheless, we note the geographic origin of several modern outbreaks of coronaviruses in East Asia, point to East Asia being a likely location where these ancient populations came into contact with the virus. Given that multiple recently recorded coronavirus outbreaks have been traced to zoonoses (direct or indirect with other animal intermediates) from East Asian bats (Wong et al., 2019), our results suggest that East Asia might have also been a natural range for coronavirus reservoir species during the last 25,000 years.

### Applied evolutionary medicine: using evolutionary information to combat COVID-19

The net result of the ancient selection patterns on the CoV-VIPs in ancient human populations is the creation of genetic differences amongst individuals now living in East Asia, and between East Asians and populations distributed across the rest of the world. As we demonstrate in this study, this evolutionary genetic information can be exploited by statistical analyses to identify loci that are potentially involved in the epidemiology of modern diseases – COVID-19 in the present case. Such evolutionary information may ultimately assist in the development of future drugs and therapies, by complementing information obtained from more traditional epidemiological and biomedical research. For example, a recent study focusing on *TMPRSS2* – a gene encoding for a transmembrane protein that plays a key role in SARS-CoV-2 infection – found that East Asian populations carry two protein coding variant that are correlated with low fatality rate for COVID-19 cases (Jeon et al., 2020). While such studies provide high quality information on a specific gene, the evolutionary approach adopted here is able to leverage evolutionary information embedded in modern genomes to identify candidate genomic regions of interest. This is similar to the information provided by GWAS – i.e. lists of variants or genes that are potentially associated with a particular trait or disease – though we note that the information provided by evolutionary analyses comes with an added understanding about the historical processes that created the underlying population genetic patterns.

The current limitation shared by population genomic approaches such as GWAS and the evolutionary analyses presented here, is that they identify statistical associations, rather than causal links, between genomic regions and traits, thereby necessitating additional research to confirm causality. In addition to the various forms of empirical information that we provide here, further evidence of causal relationships between the CoV-VIPs and COVID-19 etiology could be obtained by examining which viral proteins the selected CoV-VIPs interact with, thus establishing the specific viral functions that are affected. As a preliminary observation, we find that the 35 of the 42 selected SARS-CoV-2 VIPs tend to interact with more viral proteins than expected by chance (13 instead of six expected, see SI). Such information will help establish genetic causality and will also improve our understanding of how hosts adapt in response to viruses.

The ultimate confirmation of causality requires functional validation that the genes interact with the virus, or that drugs targeting these genes have a knock-on impact for the virus. Notably, several CoV-VIP genes are existing drug targets showing the functional importance of these particular loci (Table S7), several of which are currently being investigated or used to treat severe cases in the current COVID-19 pandemic. It remains to be established if the other genes we have identified in this study might also help guide drug repurposing efforts and provide a basis for future drug and therapeutic development to combat COVID-19 and related pathologies. It also remains to be established if population-specific past adaptation, and the underlying selected changes at those genes, could imply different drug efficacies in different human populations.

## Conclusion

By leveraging the evolutionary information contained in publicly available human genomic datasets, we were able to infer ancient viral epidemics impacting the ancestors of contemporary East Asian populations, which initially arose likely more than 20,000 years ago, resulting in coordinated adaptive changes across 42 genes. Importantly, our evolutionary genomic analyses have identified several new candidate genes that might benefit current efforts to combat COVID-19, either by providing novel drug targets or by repurposing currently available drugs that target these candidate genes (Tables S4 & S6). More broadly, our findings highlight the utility of thinking about the possible contribution of evolutionary genomic approaches into standard medical research protocols. Indeed, by revealing the identity of our ancient pathogenic foes, evolutionary genomic methods may ultimately improve our ability to predict – and thus prevent – the epidemics of the future.

## Methods

**Important note: for convenience, the 42 CoV-VIPs that we infer to have started coming under selection around 900 generations ago are called peak-VIPs in the Methods**.

### Key resources table

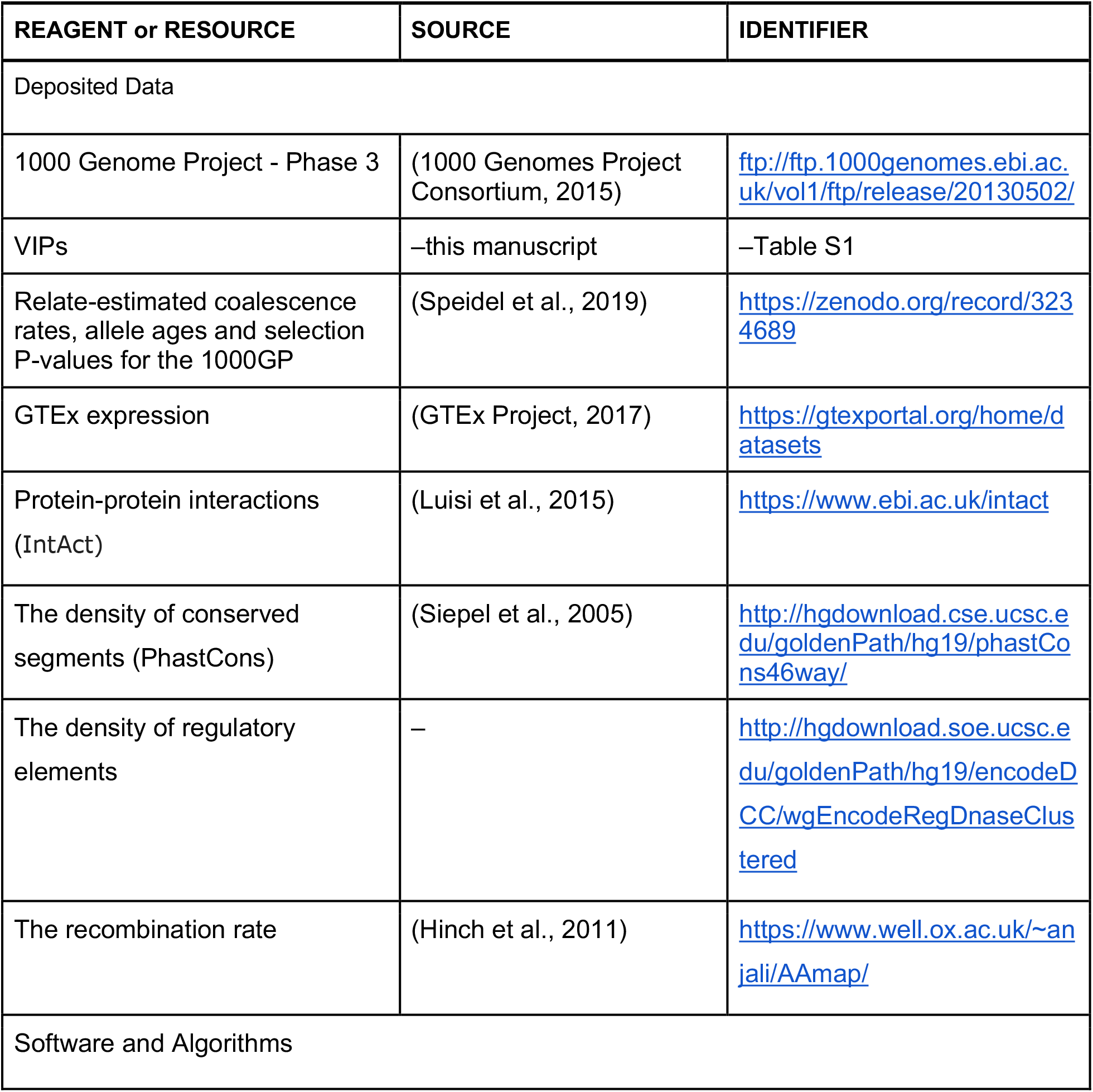
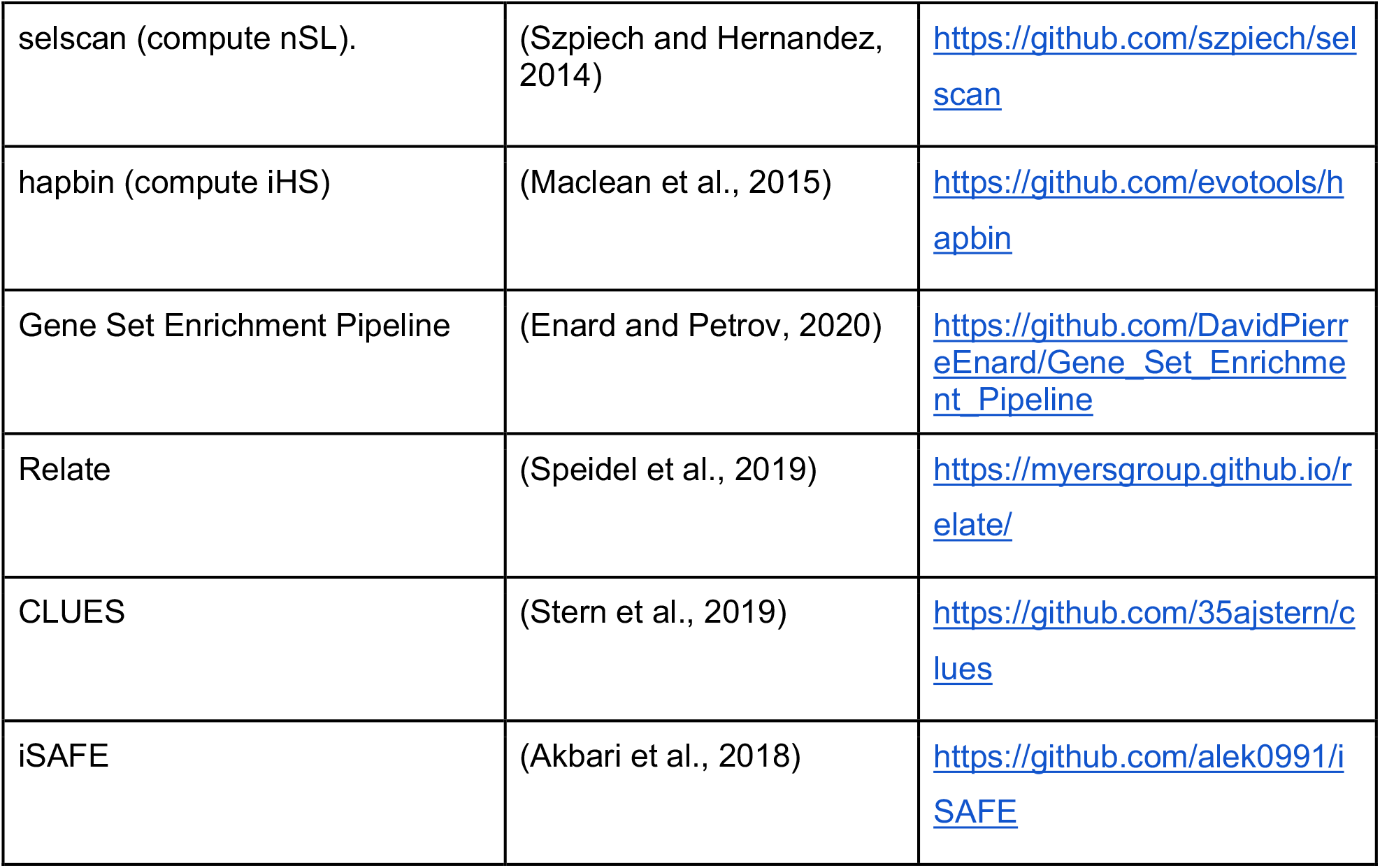

### Coronavirus VIPs

We used a dataset of 5,291 VIPs (Table S1). Of these, 1,920 of these VIPs are high confidence VIPs identified by low-throughput molecular methods, while the remaining VIPs were identified by diverse high-throughput mass-spectrometry studies. For a more detailed description of the VIPs dataset, please refer to SI: Host adaptation is expected at VIPs.

### Genomes and sweeps summary statistics

To detect signatures of adaptation in various human populations, we used the 1,000 Genome Project phase 3 dataset which provides chromosome level phased data for 26 distinct human populations representing all major continental groups (1000 Genomes Project Consortium, 2015). To measure nSL separately in each of the 26 populations, we used the selscan software available at https://github.com/szpiech/selscan (Szpiech and Hernandez, 2014). To measure iHS, we used the hapbin software available at https://github.com/evotools/hapbin (Maclean et al., 2015).

### Ranking of sweep signals at protein-coding genes and varying window sizes

To detect sweep enrichments at CoV-VIPs, we first order, separately in each of the 26 1,000 Genomes populations, human Ensembl (Cunningham et al., 2019) (version 83) protein-coding genes according to the intensity of the sweep signals at each gene. As a proxy for the intensity of these signals, we use the average of either iHS or nSL across all the SNPs with iHS or nSL values within a window of fixed size, centered at the genomic center of genes, halfway between the most upstream transcription start site and the most downstream transcription end site. We then rank the genes according to the average iHS or nSL (more precisely their absolute values) in these windows. We get six rankings for six different fixed window sizes: 50kb, 100kb, 200kb, 500kb, 1,000kb and 2,000kb. We do this to account for the variable size of sweeps of different strengths. We then estimate the sweep enrichment at CoV-VIPs compared to controls over all these different window sizes considered together, or at specific sizes, as described below and in Enard & Petrov (Enard and Petrov, 2020).

### Estimating the whole ranking curve enrichment at CoV-VIPs and its statistical significance

To estimate a sweep enrichment in a set of genes, a typical approach is to use the outlier approach to select, for example, the top 1% of genes with the most extreme signals. Here we use a previously described approach to estimate a sweep enrichment while relaxing the requirement to identify a single top set of genes. Instead of, for example, only estimating an enrichment in the top 100 genes with the strongest sweep signals, we estimate the enrichment over a wide range of top X genes, where X is allowed to vary from the top 10,000 to the top 10 with many intermediate values. This creates an enrichment curve as in Figure 1. Figure 1 shows the estimated relative fold enrichments at CoV-VIPs compared to controls, from the top 1,000 to the top 10 nSL. The statistical significance of the whole enrichment curve can then be estimated by using block-randomized genomes, as described in Enard & Petrov (Enard and Petrov, 2020). In brief, block-randomized genomes make it possible to generate a large number of random whole enrichment curves while maintaining the same level of clustering of genes in the same candidate sweeps as in the real genome, which effectively controls for gene clustering. Comparing the real whole enrichment curve to the random ones then makes it possible to estimate an unbiased false-positive risk (also known as False Discovery Rate in the context of multiple testing) for the observed whole enrichment curve at CoV-VIPs. A single false positive risk can be estimated for not just one curve but by summing over multiple curves combined, thus making it possible to estimate a single false positive risk over any arbitrary numbers of rank thresholds, window sizes, summary statistics, and populations. For instance, we estimate the false-positive enrichment risk of P=2.10^−4^ at CoV-VIPs for rank threshold from the top 10,000 to top 10, over six window sizes, for the five East Asian populations in the 1,000 Genomes data, and for both nSL and iHS, all considered together at once. This makes our approach more versatile and sensitive to selection signals ranging from a few very strong sweeps, to many, more moderately polygenic hitchhiking signals. The entire pipeline to estimate false-positive risks with block-randomized genomes is available at https://github.com/DavidPierreEnard/Gene_Set_Enrichment_Pipeline (Enard and Petrov, 2020).

### Building sets of controls matching for confounding factors

To estimate a sweep enrichment at CoV-VIPs, we compare CoV-VIPs with random control sets of genes selected far enough (>500kb) from CoV-VIPs that they are unlikely to overlap the same large sweeps. We do not compare CoV-VIPs with completely random sets of control genes. Instead, we use a previously described bootstrap test to build random control sets of genes that match CoV-VIPs for a number of potential confounding factors that might explain a sweep enrichment, rather than interactions with viruses. The bootstrap test has been described in detail (Enard and Petrov, 2020), and is available at https://github.com/DavidPierreEnard/Gene_Set_Enrichment_Pipeline.

We include 11 different potential confounding factors in the bootstrap test:

- average GTEx expression in 53 GTEx V6 tissues.
- GTEx expression in lymphocytes.
- GTEx expression in testis.
- the number of protein-protein interactions from the Intact database, curated by Luisi et al. (Luisi et al., 2015).
- the Ensembl (v83) coding sequence density in a 50kb window centered on each gene.
- the density of conserved segments identified by PhastCons (Siepel et al., 2005) (http://hgdownload.cse.ucsc.edu/goldenPath/hg19/phastCons46way/).
- the density of regulatory elements, estimated by the density of Encode DNase I V3 Clusters (http://hgdownload.soe.ucsc.edu/goldenPath/hg19/encodeDCC/wgEncodeRegDnaseClustered/) in a 50kb window centered on each gene.
- the recombination rate in a 200kb window centered on each gene (Hinch et al., 2011).
- the GC content in a 50kb window centered on each gene.
- the number of bacteria each gene interacts with, according to the Intact database (as of June 2019; https://www.ebi.ac.uk/intact/).
- the proportion of genes that are immune genes according to Gene Ontology annotations GO:0006952 (defense response), GO:0006955 (immune response), and GO:0002376 (immune system process) as of May 2020.

### Estimating adaptation start times at specific genes with Relate

As times of emergence of adaptive mutations, we use the publicly available estimates from Relate (https://myersgroup.github.io/relate/). Relate estimates mutation emergence times while controlling for fluctuations of population size over time, based on the coalescence rates it reconstructs after inferring ancestral recombination graphs at the scale of the whole genome (Speidel et al., 2019). Relate provides two times of emergence of mutations, one low estimate (less generations ago), and one high estimate (more generations ago). The low time estimate corresponds to the time when Relate estimates an elevated probability that the frequency of the mutation is different from zero. The high time estimate corresponds to the time when Relate estimates that the probability is not too small that the frequency of the mutation is different from zero. For our purpose of estimating when selection started, the low time estimate is the best suited, because it provides an estimate of when the frequency of a selected mutation was already high enough to distinguish from zero, for those mutations where selection started from a very low frequency. For cases where selection started with standing genetic variants that were already distinguishable from zero, the Relate low time estimates for the emergence of mutations do not provide a good proxy for when selection actually started. Thus, if we were able to estimate when selection started for standing genetic variants, we might be able to observe an even stronger peak than the one we see when just relying on those variants where selection started from low frequencies.

Using the low Relate time estimates is also justified due to the fact that the sweep establishment phase can take very variable amounts of time before the start of the sweep exponential phase. During the establishment phase, selected alleles are still mostly governed by drift which makes pinpointing the actual starting time of selection difficult. In this context, the low Relate time estimates provide an estimate of the time when the selected alleles were no longer at very low frequencies not statistically different from zero, and closer to entering the exponential phase, which provides a more certain time estimate for when selection started for certain.

An important step is then to choose at each CoV-VIP locus, and all the other control loci, which Relate mutation to use to get a single time estimate for each locus. Note that here we make an assumption that each locus has experienced only one single adaptive event. Given our finding that iSAFE peaks at CoV-VIPs are much closer to GTEx V8 eQTLs than expected by chance, it is likely that the selected adaptive mutations are regulatory mutations at, or close to annotated eQTLs for a specific gene. They are not necessarily exactly located at eQTLs, because current eQTLs annotations may still be incomplete, and in our case we use eQTLs identified in GTEx V8 using mostly European individuals, even though we analyse selection signals in East Asian populations. Because of these limitations, we use the Relate estimated time at the mutation where Relate estimates the lowest positive selection p-value within 50kb windows centered on eQTLs. We also only consider variants with a minor allele frequency greater than 20%, given the signals detected by iHS and nSL that only have some power to detect incomplete sweeps above 20% frequencies (Ferrer-Admetlla et al., 2014; Voight et al., 2006). This also excludes a potential risk of confounding by low frequency neutral or weakly deleterious variants, that can show selection-like patterns when their only way to escape removal early on is through a chance, rapid frequency increase that can look like selection. The Relate selection test is based on faster than expected coalescence rates given the population size at any given time, and its results are publicly available at https://myersgroup.github.io/relate/. Note that the mutation with the lowest Relate p-value does not always overlap with an iSAFE peak (Figure S5), which is not entirely surprising if the haplotype signals exploited by both Relate and iSAFE partly deteriorated due to recombination since the time selection at CoV-VIPs was strong (Figures 3 and 5). Both of these methods are indeed designed to locate the selected variant right after, or during, active selection.

Because we work with five different East Asian populations, we more specifically select the variant with the lowest Relate selection test p-value on average across all the five East Asian populations. Then, we also use the corresponding average low Relate mutation time estimate across the five East Asian populations. We do not attempt to estimate the selection time and p-value by considering all 1,000 Genomes East Asian individuals tested together by Relate, because then the Relate selection test is at a greater risk of being confounded by population structure. Finally, we only consider CoV-VIPs and other control genes with an average Relate selection test p-value lower than 10^−3^, to make sure that we indeed use estimated times at selected variants.

### The peak significance test

To test if the peak of Relate time estimates around 900 generations ago at CoV-VIPs (Figure 2) is expected simply by chance or not, we designed a peak significance test. The test compares the peak at CoV-VIPs, with the top peaks obtained when repeatedly randomly sampling sets of genes. We first identify the most prominent peak at CoV-VIPs by visual inspection of the pink distribution of Relate times for CoV-VIPs compared to the blue distribution of Relate times for all protein-coding genes with an estimated Relate time (Figure 2). To build these distributions, top Relate selected mutations shared between multiple neighboring genes (CoV-VIPs or controls) are counted only once, to avoid a confounding effect of gene clustering (152 selected variants at CoV-VIPs, 1771 selected variants for all protein coding genes). The peak around 900 generations ago (870 generations more exactly) spans approximately 200 generations, where the pink distribution is clearly above the blue one. We then use a 200 generations-wide window, sliding every generation from 0 to 6,000 generations to verify the peak more rigorously. Sliding one generation after another, each time we count the difference between the number of Relate selected variants at CoV-VIPs that fall in the sliding 200 generations window, and the number of Relate selected variants at all other genes that are not CoV-VIPs, weighted by the percentage of variants found at CoV-VIPs, to correct for the different size of the two sets of variants. Using this sliding window approach, the top of the peak is found at 870 generations, with a difference of 19.5 additional Relate selected variants between 770 and 970 (870 plus or minus 100) at CoV-VIPs compared to the null expectation.

We then repeat the sliding of a 200 generations window to identify the maximum peak and measure the same difference, but this time for random sets of Relate selected variants of the same size (152 selected variants out of the 1,771 selected variants). To estimate p-values, we then compare the actual observed difference with the distribution of differences generated with one million random samples.

As mentioned in the Results, one potential issue is that we run the peak significance test after we already know that CoV-VIPs are enriched for iHS and nSL top sweeps, and especially enriched for nSL top sweeps. This enrichment may skew the null expectation for the distribution of Relate times at CoV-VIPs. In other words, there is a risk that any set of genes with the same sweep enrichment might exhibit the same peak as CoV-VIP. As a result, comparing CoV-VIPs with randomly chosen non-CoV-VIPs may not be appropriate. To test this, we repeat the peak significance test, but this time comparing the peak at CoV-VIPs with the peaks at random sets of non-CoV-VIPs that we build to have the same distribution of nSL ranks as CoV-VIPs. To do this, we define nSL bins between ranks 1 and the highest rank with a rank step of 100 between each bin, and we count how many Relate selected variants fall in each bin (each gene has one nSL rank and one Relate selected variant). To build the random set, we then fill each of the 100 bins with the same number of random non-CoV-VIPs, as long as their nSL rank falls within that bin. We use the average nSL rank over the five East Asian populations, and the lower population-averaged rank of either 1 Mb or 2Mb window sizes (where we observe the strongest enrichment at CoV-VIPs, see Results). The results of the peak significance test are unchanged when using the matching nSL distribution (peak significance test P=1.10-4 vs. P=2.3.10-4 without matching nSL distribution).

In further agreement with the fact that the sweep enrichment does not confound the peak significance test, the peak at CoV-VIPs stands out more when repeating the peak significance test using a smaller nSL top rank limit (Figure S6). In this case, we compare sets of CoV-VIPs and sets of controls both enriched in stronger sweep signals. Thus, if stronger sweep signals at CoV-VIPs biased the peak significance test, we would expect the peak to fade away when comparing only CoV-VIPs and controls both with stronger nSL signals. Conversely, we observe that half of the CoV-VIPs with the weaker nSL signals (population-averaged nSL rank higher than 7,200 for both 1Mb and 2Mb windows) do not show a significant peak (peak significance test P=0.53).

### The iSAFE peaks/eQTL proximity test

Adaptation in the human genome was likely mostly regulatory adaptation through gene expression changes (Enard et al., 2014; Kudaravalli et al., 2009; Nédélec et al., 2016; Quach et al., 2016). To test if positive selection at CoV-VIPs likely involved regulatory changes, we ask whether the signals of adaptation around CoV-VIPs are localized closer than expected by chance to GTEx eQTLs that affect the expression of CoV-VIPs in present human populations. Indeed, the genomic regions at or close to CoV-VIP GTEx eQTLs are likely enriched for CoV-VIP regulatory elements, and therefore the most likely place to find CoV-VIP-related adaptations in the genome. To localize where adaptation occurred, we use the iSAFE method that was specifically designed for this purpose (Akbari et al., 2018). iSAFE scans the genome and estimates a score that increases together with proximity to the actual selected mutation. The higher the score, the higher the odds that the scored variant is itself the selected one, or close to the selected one. An important caveat is that iSAFE is designed to localize where selection happened right after it happened, or as selection is still ongoing. In our case, we have evidence that selection was strong at CoV-VIPs only more than 500 generations (∼14,000 years) ago, and then much weaker more recently (Figure 5). This could be an issue, because we expect that recombination events that occurred after the strong selection might have deteriorated the iSAFE signal that relies on haplotype structure. This is because recombination mixes together the haplotypes that hitchhiked with the selected mutation, with those that did not. In line with this, we often do not observe simple, clean iSAFE score peaks, but instead, iSAFE score plateaus and more rugged peaks (Figure S5). For this reason, we designed an approach to not only identify the top of simple iSAFE peaks, but also more rugged peaks or plateaus. First, to measure iSAFE scores, we combine all the haplotypes from the five East Asian populations together as input, since we found that the selection signal at CoV-VIPs is common to all these populations (iSAFE parameters: −-IgnoreGaps --MaxRegionSize 250000000 --window 300 -- step 100 --MaxFreq .95 --MaxRank 15). We then use a 500kb window sliding every 10kb to identify the highest local iSAFE value in the 500kb window (Figure S8). Once we have the highest local iSAFE value and coordinate, we define a broader iSAFE peak as the region both upstream and downstream where the iSAFE values are still within 80% of the maximum value (Figure S8). This way, we can better annotate iSAFE plateaus and rugged peaks, and take into account the fact that they can span more than just a narrow local maximum (Figure S5).

Once the local iSAFE peaks are identified, we can ask how close GTEx eQTLs are to these peaks compared to random expectations. We first measure the distance of each CoV-VIP GTEx eQTL to the closest iSAFE peak. To avoid redundancy, we merge eQTLs closer than 1kb to each other into one test eQTL at the closest, lower multiple of 1,000 genomic coordinates (for example 3,230 and 3,950 would both become 3,000). We then measure the average of the log of the distance between all CoV-VIPs and their closest iSAFE peak. We use the log (base 10) of the distance, because it matters if the eQTL/iSAFE peak distance is 100 bases instead of 200kb, but it does not really matter if the distance is 200kb or 600kb, because the iSAFE peak at 300kb is likely not related to the eQTL more than the peak at 600kb. Once we have the average of log-distances, we compare it to its random expected distribution. To get this random distribution, we measure the log-distance between each CoV-VIP eQTL and the iSAFE peaks, but after shifting the iSAFE scores left or right by a random value between 1Mb and 2Mb (Figure S8; less, or no shift at all if this falls within telomeres or centromeres). We shift by at least 1Mb to make sure that we do not rebuild the original overlap of iSAFE peaks with eQTLs again and again (some iSAFE peaks, or more precisely rugged peaks and plateaus can be wide and include several hundred kilobases; see Figure S5). The random shifting effectively breaks the relationship between eQTLs and iSAFE peaks, while maintaining the same overall eQTL and peak structure (and thus variance for the test). The random log-distance distribution then provides an overall random average log-distance to compare the observed average long-distance with, as well as estimate a p-value.

Then, to more specifically ask if lung eQTLs at CoV-VIPs or the eqTLs of other specific tissues are closer to iSAFE peaks than expected by chance, we can do the same but only using the eQTLs of that specific tissue. The analysis represented in figure 6 is however more complicated than just testing if CoV-VIP eQTLs for a specific tissue are closer to iSAFE peaks than expected by chance by randomly sliding iSAFE values. Instead, what we ask is whether the 42 peak-VIPs have eQTLs for a given tissue that are even closer to iSAFE peaks than the eQTLs of all CoV-VIPs in general. To test this, for example with lung eQTLs, we first estimate how close lung eQTLs are to iSAFE peaks at peak-VIPs, compared to random expectations, by measuring the difference between the observed and the average random log-distance, just as described before. We then count the number of peak-VIPs with lung eQTLs (19 out of 25 peak-VIPs with GTEx eQTLs), and we randomly select the same number of any CoV-VIP (which may randomly include peak-VIPs) as long as the random set of CoV-VIPs has the same number of lung eQTLs (plus or minus 10%) as the set of peak VIPs with lung eQTLs (the same gene can have multiple eQTLs for one tissue). We make sure that the tested and the random sets have similar numbers of genes and eQTLs so that the test has the appropriate null variance. We then measure the difference between the observed log-distance, and the randomly expected average log-distance for the random set of CoV-VIPs, exactly the same way we did before for the actual set of peak-VIPs. We then measure the ratio of the observed difference in log-distance between peak-VIPs and the random expectation after many random shiftings (1,000), divided by the average of the same difference measured over many random sets of CoV-VIPs. The final ratio tells us how much closer lung eQTLs are to iSAFE peaks at peak-VIPs compared to CoV-VIPs in general, and still takes the specific eQTLs and iSAFE peak structures at each locus into account, since we compare differences in log-distances expected while preserving the same eQTL and iSAFE peak structure (see above the description of the random coordinate shifting). One important last detail about the test is that because we already found that the 50% of loci with the lowest nSL signals do not show a peak of selection at CoV-VIPs around 900 generations ago (see Results), we do not use these loci in this test since any iSAFE peak there is much more likely to represent random noise, not actual selection locations, and thus likely to dilute genuine signals. Using this test, we find that lung and other tissues’ eQTLs at peak-VIPs are much closer to iSAFE peaks than they are at CoV-VIPs in general. This test thus specifically tells that adaptation happened closer to lung eQTLs, specifically around 900 generations ago compared to other evolutionary times. By estimating the same ratio for 24 other tissues with at least 10 peak-VIPs with the specific tested tissue eQTLs, we can finally rank each tissue for its more pronounced involvement in adaptation ∼900 generations ago, as done in figure 6. It is particularly interesting in this respect that the tissue with least evidence for being more involved in adaptation at that time more than other evolutionary times is spleen. Spleen indeed likely represents a good negative control as a tissue strongly enriched in immune cell types and likely to have evolved adaptively for most of evolution.

### UK Biobank GWAS analysis

To compare the UK Biobank GWAS p-values at different loci, we assigned one p-value for each gene, either CoV-VIPs, peak-VIPs or other genes, even though each gene locus can have many variants with associated GWAS p-values. To assign just one single GWAS p-value to each gene, we selected the variant with the lowest p-value at or very close (<1kb) to GTEx eQTLs for a specific gene, in line with the fact that GWAS hits tend to overlap eQTLs (Hormozdiari et al., 2016), and to remain consistent with the rest of our manuscript. We then compared the average p-value between different sets of genes using classic permutations (one billion iterations).

### Drug targets identification

We queried the databases DGIdb (Cotto et al., 2017), and PanDrugs (Piñeiro-Yáñez et al., 2018) for drugs targeting CoV-VIPs and peak-VIPs. For hits from PanDrugs we limited the results to only genes that are in direct interaction with the designated drug. Drugs targeting peak-VIPs are presented in Table S7. In addition, we present a list of peak-VIPs that are not currently drug targets, but have been previously identified in (Finan et al., 2017) as viable drug targets (druggable genome).

## Acknowledgments

We wish to thank Leo Speidel and Aaron Stern for their valuable help using Relate and CLUES, respectively. Y.S. is supported by the Australian Research Council (ARC DP190103705). R.T. is an ARC DECRA fellow (DE190101069).

## Authors Contributions

Conceived and designed the experiments: YS, RT, DE. Performed the experiments: YS, MEL, RT, DE. Interpreted the results: YS, MEL, RT,CDH, ASJ, DE. Wrote the manuscript: YS, RT, DE

## Notes

### Competing Interest Statement

The authors have declared no competing interest.

### Summary of Updates

We revised the title; WE addressed more directly multiple misconceptions about the results that arose on social media; We corrected a mis-specified parameter when using CLUES (the time cutoff tCutOff, pushed from 1,000 to 1,500 generations) to allow inference of older non-zero allele frequencies and Figure 3 has been revised accordingly; We added the frequency trajectories estimated by CLUES in the Yoruba and in the British populations; We added a supplemental table with the current frequencies of the selected alleles in Africa, Europe and East Asia, together with the corresponding SNP IDs; We added justifications about why we use the UK Biobank COVID-19 GWAS instead of other alternatives to avoid population stratification issues as much as possible.

## References

1000 Genomes Project Consortium (2015). A global reference for human genetic variation. Nature 526, 68–74.

Akbari, A., Vitti, J.J., Iranmehr, A., Bakhtiari, M., Sabeti, P.C., Mirarab, S., and Bafna, V. (2018). Identifying the favored mutation in a positive selective sweep. Nat. Methods 15, 279–282.

Balogun, O.D., Bea, V.J., and Phillips, E. (2020). Disparities in Cancer Outcomes Due to COVID-19—A Tale of 2 Cities. JAMA Oncol.

Bao, C., Tao, X., Cui, W., Yi, B., Pan, T., Young, K.H., and Qian, W. (2020). SARS-CoV-2 induced thrombocytopenia as an important biomarker significantly correlated with abnormal coagulation function, increased intravascular blood clot risk and mortality in COVID-19 patients. Exp. Hematol. Oncol. 9, 16.

Barreiro, L.B., Ben-Ali, M., Quach, H., Laval, G., Patin, E., Pickrell, J.K., Bouchier, C., Tichit, M., Neyrolles, O., Gicquel, B., et al. (2009). Evolutionary dynamics of human Toll-like receptors and their different contributions to host defense. PLoS Genet. 5, e1000562.

Best, S.M., and Kerr, P.J. (2000). Coevolution of host and virus: the pathogenesis of virulent and attenuated strains of myxoma virus in resistant and susceptible European rabbits. Virology 267, 36–48.

Cotto, K.C., Wagner, A.H., Feng, Y.-Y., Kiwala, S., Coffman, A.C., Spies, G., Wollam, A., Spies, N.C., Griffith, O.L., and Griffith, M. (2017). DGIdb 3.0: a redesign and expansion of the drug– gene interaction database. Nucleic Acids Res. 46, D1068–D1073.

Cunningham, F., Achuthan, P., Akanni, W., Allen, J., Amode, M.R., Armean, I.M., Bennett, R., Bhai, J., Billis, K., Boddu, S., et al. (2019). Ensembl 2019. Nucleic Acids Res. 47, D745–D751.

Dong, E., Du, H., and Gardner, L. (2020). An interactive web-based dashboard to track COVID-19 in real time. Lancet Infect. Dis. 20, 533–534.

eGTEx Project (2017). Enhancing GTEx by bridging the gaps between genotype, gene expression, and disease. Nat. Genet. 49, 1664–1670.

Ellinghaus, D., Degenhardt, F., Bujanda, L., Buti, M., Albillos, A., Invernizzi, P., Fernández, J., Prati, D., Baselli, G., Asselta, R., et al. (2020). Genomewide Association Study of Severe Covid-19 with Respiratory Failure. N. Engl. J. Med.

Elmunzer, B.J., Spitzer, R.L., Foster, L.D., Merchant, A.A., Howard, E.F., Patel, V.A., West, M.K., Qayed, E., Nustas, R., Zakaria, A., et al. (2020). Digestive Manifestations in Patients Hospitalized with COVID-19. Clin. Gastroenterol. Hepatol.

Enard, D., and Petrov, D.A. (2018). Evidence that RNA Viruses Drove Adaptive Introgression between Neanderthals and Modern Humans. Cell 175, 360–371.e13.

Enard, D., and Petrov, D.A. (2020). Ancient RNA virus epidemics through the lens of recent adaptation in human genomes. Philos. Trans. R. Soc. Lond. B Biol. Sci. 375, 20190575.

Enard, D., Messer, P.W., and Petrov, D.A. (2014). Genome-wide signals of positive selection in human evolution. Genome Res. 24, 885–895.

Enard, D., Cai, L., Gwennap, C., and Petrov, D.A. (2016). Viruses are a dominant driver of protein adaptation in mammals. Elife 5, 56.

Ferrer-Admetlla, A., Liang, M., Korneliussen, T., and Nielsen, R. (2014). On detecting incomplete soft or hard selective sweeps using haplotype structure. Mol. Biol. Evol. 31, 1275– 1291.

Finan, C., Gaulton, A., Kruger, F.A., Lumbers, R.T., Shah, T., Engmann, J., Galver, L., Kelley, R., Karlsson, A., Santos, R., et al. (2017). The druggable genome and support for target identification and validation in drug development. Sci. Transl. Med. 9.

Gene Ontology Consortium (2015). Gene Ontology Consortium: going forward. Nucleic Acids Res. 43, D1049–D1056.

Gordon, D.E., Jang, G.M., Bouhaddou, M., Xu, J., Obernier, K., White, K.M., O’Meara, M.J., Rezelj, V.V., Guo, J.Z., Swaney, D.L., et al. (2020). A SARS-CoV-2 protein interaction map reveals targets for drug repurposing. Nature 583, 459–468.

Grosse, C., Grosse, A., Salzer, H.J.F., Dünser, M.W., Motz, R., and Langer, R. (2020). Analysis of cardiopulmonary findings in COVID-19 fatalities: High incidence of pulmonary artery thrombi and acute suppurative bronchopneumonia. Cardiovasc. Pathol. 49, 107263.

Hayward, L.K., and Sella, G. (2019). Polygenic adaptation after a sudden change in environment.

Hinch, A.G., Tandon, A., Patterson, N., Song, Y., Rohland, N., Palmer, C.D., Chen, G.K., Wang, K., Buxbaum, S.G., Akylbekova, E.L., et al. (2011). The landscape of recombination in African Americans. Nature 476, 170–175.

Hoffmann, C., and Kamps, B.S. (2003). SARS Reference (Flying Publisher).

Hormozdiari, F., van de Bunt, M., Segrè, A.V., Li, X., Joo, J.W.J., Bilow, M., Sul, J.H., Sankararaman, S., Pasaniuc, B., and Eskin, E. (2016). Colocalization of GWAS and eQTL Signals Detects Target Genes. Am. J. Hum. Genet. 99, 1245–1260.

Jeon, S., Blazyte, A., Yoon, C., Ryu, H., Jeon, Y., Bhak, Y., Bolser, D., Manica, A., Shin, E.-S., Cho, Y.S., et al. (2020). Ethnicity-dependent allele frequencies are correlated with COVID-19 case fatality rate (Authorea, Inc.).

Kudaravalli, S., Veyrieras, J.-B., Stranger, B.E., Dermitzakis, E.T., and Pritchard, J.K. (2009). Gene expression levels are a target of recent natural selection in the human genome. Mol. Biol. Evol. 26, 649–658.

Luisi, P., Alvarez-Ponce, D., Pybus, M., Fares, M.A., Bertranpetit, J., and Laayouni, H. (2015). Recent positive selection has acted on genes encoding proteins with more interactions within the whole human interactome. Genome Biol. Evol. 7, 1141–1154.

Maclean, C.A., Chue Hong, N.P., and Prendergast, J.G.D. (2015). hapbin: An Efficient Program for Performing Haplotype-Based Scans for Positive Selection in Large Genomic Datasets. Mol. Biol. Evol. 32, 3027–3029.

Michalakis, K., and Ilias, I. (2020). SARS-CoV-2 infection and obesity: Common inflammatory and metabolic aspects. Diabetes Metab. Syndr. 14, 469–471.

Moorjani, P., Sankararaman, S., Fu, Q., Przeworski, M., Patterson, N., and Reich, D. (2016). A genetic method for dating ancient genomes provides a direct estimate of human generation interval in the last 45,000 years. Proc. Natl. Acad. Sci. U. S. A. 113, 5652–5657.

Nédélec, Y., Sanz, J., Baharian, G., Szpiech, Z.A., Pacis, A., Dumaine, A., Grenier, J.-C., Freiman, A., Sams, A.J., Hebert, S., et al. (2016). Genetic Ancestry and Natural Selection Drive Population Differences in Immune Responses to Pathogens. Cell 167, 657–669.e21.

Ou, X., Liu, Y., Lei, X., Li, P., Mi, D., Ren, L., Guo, L., Guo, R., Chen, T., Hu, J., et al. (2020). Characterization of spike glycoprotein of SARS-CoV-2 on virus entry and its immune cross-reactivity with SARS-CoV. Nat. Commun. 11, 1620.

Piñeiro-Yáñez, E., Reboiro-Jato, M., Gómez-López, G., Perales-Patón, J., Troulé, K., Rodríguez, J.M., Tejero, H., Shimamura, T., López-Casas, P.P., Carretero, J., et al. (2018). PanDrugs: a novel method to prioritize anticancer drug treatments according to individual genomic data. Genome Med. 10, 41.

Quach, H., Rotival, M., Pothlichet, J., Loh, Y.-H.E., Dannemann, M., Zidane, N., Laval, G., Patin, E., Harmant, C., Lopez, M., et al. (2016). Genetic Adaptation and Neandertal Admixture Shaped the Immune System of Human Populations. Cell 167, 643–656.e17.

Quintana-Murci, L. (2019). Human Immunology through the Lens of Evolutionary Genetics. Cell 177, 184–199.

Richman, D.D., Whitley, R.J., and Hayden, F.G. (2020). Clinical Virology (John Wiley & Sons).

Roberts, G.H.L., Park, D.S., Coignet, M.V., McCurdy, S.R., Knight, S.C., Partha, R., Rhead, B., Zhang, M., Berkowitz, N., Baltzell, A.K.H., et al. (2020). AncestryDNA COVID-19 Host Genetic Study Identifies Three Novel Loci. medRxiv.

Sabeti, P.C., Schaffner, S.F., Fry, B., Lohmueller, J., Varilly, P., Shamovsky, O., Palma, A., Mikkelsen, T.S., Altshuler, D., and Lander, E.S. (2006). Positive natural selection in the human lineage. Science 312, 1614–1620.

Sattar Naveed, McInnes Iain B., and McMurray John J.V. (2020). Obesity Is a Risk Factor for Severe COVID-19 Infection. Circulation 142, 4–6.

Sawyer, S.L., Wu, L.I., Emerman, M., and Malik, H.S. (2005). Positive selection of primate TRIM5α identifies a critical species-specific retroviral restriction domain. Proc. Natl. Acad. Sci. U. S. A. 102, 2832–2837.

Scarpone, C., Brinkmann, S.T., Große, T., Sonnenwald, D., Fuchs, M., and Walker, B.B. (2020). A multimethod approach for county-scale geospatial analysis of emerging infectious diseases: a cross-sectional case study of COVID-19 incidence in Germany. Int. J. Health Geogr. 19, 32.

Schrider, D.R. (2020). Background Selection Does Not Mimic the Patterns of Genetic Diversity Produced by Selective Sweeps. Genetics 216, 499–519.

Siepel, A., Bejerano, G., Pedersen, J.S., Hinrichs, A.S., Hou, M., Rosenbloom, K., Clawson, H., Spieth, J., Hillier, L.W., Richards, S., et al. (2005). Evolutionarily conserved elements in vertebrate, insect, worm, and yeast genomes. Genome Res. 15, 1034–1050.

Speidel, L., Forest, M., Shi, S., and Myers, S.R. (2019). A method for genome-wide genealogy estimation for thousands of samples. Nat. Genet. 51, 1321–1329.

Stern, A.J., Wilton, P.R., and Nielsen, R. (2019). An approximate full-likelihood method for inferring selection and allele frequency trajectories from DNA sequence data. PLoS Genet. 15, e1008384.

Stern, A.J., Speidel, L., Zaitlen, N.A., and Nielsen, R. (2020). Disentangling selection on genetically correlated polygenic traits using whole-genome genealogies. bioRxiv.

Sudlow, C., Gallacher, J., Allen, N., Beral, V., Burton, P., Danesh, J., Downey, P., Elliott, P., Green, J., Landray, M., et al. (2015). UK biobank: an open access resource for identifying the causes of a wide range of complex diseases of middle and old age. PLoS Med. 12, e1001779.

Szpiech, Z.A., and Hernandez, R.D. (2014). selscan: an efficient multithreaded program to perform EHH-based scans for positive selection. Mol. Biol. Evol. 31, 2824–2827.

Tajima, F. (1989). Statistical method for testing the neutral mutation hypothesis by DNA polymorphism. Genetics 123, 585–595.

The COVID-19 Host Genetics Initiative (2020). The COVID-19 Host Genetics Initiative, a global initiative to elucidate the role of host genetic factors in susceptibility and severity of the SARS-CoV-2 virus pandemic. Eur. J. Hum. Genet. 28, 715–718.

Uricchio, L.H., Petrov, D.A., and Enard, D. (2019). Exploiting selection at linked sites to infer the rate and strength of adaptation. Nat Ecol Evol 3, 977–984.

Voight, B.F., Kudaravalli, S., Wen, X., and Pritchard, J.K. (2006). A map of recent positive selection in the human genome. PLoS Biol. 4, e72.

Wong, A.C.P., Li, X., Lau, S.K.P., and Woo, P.C.Y. (2019). Global Epidemiology of Bat Coronaviruses. Viruses 11.

World Health Organization (2019). Middle East respiratory syndrome coronavirus (MERS-CoV).

Zeberg, H., and Pääbo, S. (2020). The major genetic risk factor for severe COVID-19 is inherited from Neandertals.

