## Supplemental Information for "An ancient viral epidemic involving host coronavirus interacting genes more than 20,000 years ago in East Asia"

### Supplemental analyses

For convenience, the 42 CoV-VIPs with selection starting around 900 generations ago are called peak-VIPs.

### Host adaptation is expected at VIPs.

Using VIPs to find the genomic footprints of an ancient epidemic is justified by the fact that VIPs do not just interact with viruses. These interactions are in fact functionally consequential for viruses. The 420 CoV-VIPs are part of a much larger set of VIPs found to interact to date with more than 20 different viruses that infect humans (Enard and Petrov, 2018). In total, there are currently 5,291 VIPs (Table S1). Of these, 1,920 high confidence VIPs were annotated manually by curating the virology literature and corresponding to VIPs that were identified by low-throughput molecular methods (Enard and Petrov, 2018). These VIPs were often identified by virologists who hypothesized that the interaction existed in the first place based on previous virology knowledge. The other 3,371 VIPs identified by multiple high-throughput mass spectrometry experiments, such as the one conducted to identify the 332 SARS-CoV-2 VIPs (Gordon et al., 2020).

To confirm that VIPs are indeed functionally important for viruses beyond just interacting physically, and represent a viable way of detecting specific viral selective pressures that trigger host adaptation, we verify that VIPs have antiviral or proviral effects on the viral replication cycle on which positive selection can act. More specifically, we need to confirm that VIPs have much more frequent proviral or antiviral effects compared to non-VIPs. To test this, we are currently manually annotating all protein-coding genes in the human genome that were involved in published low-throughput expression perturbation experiments to assess their effects on viruses, and manually curable in PubMed. Such expression perturbation experiments typically include RNAi knock-down experiments or overexpression experiments. These experiments are useful to annotate proviral or antiviral effects. Indeed, decreasing the expression of an antiviral VIP should be beneficial to viral replication, while increasing the expression of an antiviral should be detrimental to the virus. Conversely, decreasing the expression of a proviral VIP should be detrimental to viral replication, while increasing the expression of a proviral VIP should be beneficial. We consider only low-throughput expression perturbation experiments, where the expression of only one candidate gene is perturbed. This excludes high throughput genome-wide RNAi screens notorious for their high false positive and high false negative rates. Using these criteria, we have so far found that 855, or 66% of 1,300 already annotated low-throughput VIPs have a known antiviral or proviral effect. Of the 2,627 high-throughput VIPs that we already annotated, 426 or 16% have a known antiviral or proviral effect. Of the 3,913 non-VIPs that we already annotated, 171 or 4% have a known antiviral of proviral effect. Although we have not annotated all human protein-coding genes yet, the large numbers already annotated imply that these proportions are very likely to be close to the final proportions when all genes are annotated.

Thus, approximately two-thirds of low-throughput VIPs have known antiviral or proviral effects that were revealed by expression perturbation experiments such as gene knock-down or over-expression. The 16% proportion of high throughput VIPs known to have a clear antiviral or proviral effect is much lower than the two-thirds of low-throughput VIPs with antiviral or proviral effects, but it is important to consider that high-throughput VIPs have not been investigated anywhere near as much as the low-throughput ones. In contrast, only 4% of non-VIPs with no known viral interaction have published antiviral or proviral effects. Both low-throughput and high-throughput VIPs are thus far more often functionally consequential for viruses compared to non-VIPs (simple permutation test P<10-16 in both cases). Note that because they will dilute the signal rather than create it, a certain amount of random, false-positive high-throughput interactions are expected to be conservative when trying to detect ancient epidemics.

Focusing specifically on the 420 CoV-VIPs, we find that 121 or 28.9% of them already have published antiviral or proviral effects (Table S1). Of the 332 SARS-CoV-2 VIPs, 83 or 25% of them have antiviral or proviral effects (Table S1), often independently confirmed in multiple viruses. The SARS-CoV-2 VIPs are thus more than six times more likely to have antiviral or proviral effects than non-VIPs, which supports the high quality of the mass spectrometry screen conducted by Gordon et.al. (Gordon et. al. 2020). Note that it is unrealistic to expect much higher percentages at SARS-CoV-2 VIPs, given that coronaviruses are only starting to be more thoroughly investigated, and have been much less investigated than other viruses such as HIV or IAV (influenza).

### Host intrinsic functions do not explain the pattern and timing of adaptation at CoV-VIPs.

An important limitation to consider when inferring ancient epidemics is that VIPs do not just interact with viruses, but are also involved in multiple hosts intrinsic functions. These host functions could explain the enrichment and timing of adaptation at CoV-VIPs, rather than interactions with a coronavirus or related virus. This would happen as a result of specific host functions being enriched at CoV-VIPs, and also intrinsically enriched in adaptive signals independently of any interaction with viruses. Host functions not enriched at CoV-VIPs are not expected to generate an enrichment in adaptation at CoV-VIPs, because the lack of enrichment means that they are present in similar or smaller proportions in the rest of the genome.

Thus, if host functions enriched at CoV-VIPs, rather than viral interactions, explain adaptation at CoV-VIPs, we expect that i) genes with these host functions should be enriched in sweep signals even when they don't interact with coronaviruses and ii) genes with these host functions should have started adapting around 900 generations ago, to also explain the timing of adaptation at CoV-VIPs, even when they do not interact with coronaviruses.

To estimate the role of host intrinsic functions, we use the functional annotations from the Gene Ontology (GO) for GO biological processes, GO molecular functions, and GO cellular localizations. In total, there are 106 GO annotations that are enriched at CoV-VIPs compared to the matched controls already used to assess the sweep enrichment (P<0.001 based on 10,000 matched control sets). Of these 106 GO annotations, only 20 have a more than two-fold enrichment among CoV-VIPs (and 50 genes or more among non-CoV-VIPs; Table S2) and are thus more likely to contribute to the strong sweeps enrichment at CoV-VIPs. We first test if these 20 GO annotations are enriched in sweeps independently of any interaction with coronaviruses. To do this, we use the same bootstrap test used to compare CoV-VIPs with matched controls, but this time we compare genes with the GO annotations, with control genes far enough (>500kb) from any other gene with these annotations. To make sure that a significant enrichment would have nothing to do with coronaviruses, we exclude from this comparison any gene closer than 500kb to any CoV-VIP. In total, there are 1723 genes with at least one of the 20 highly enriched GO annotations, and 3701 far enough potential control genes. Using exactly the same iHS and nSL enrichment curves used to detect a sweep enrichment at CoV-VIPs (STAR Methods), we do not find any significant enrichment at the 1723 genes compared to matched controls (whole enrichment curves for nSL and iHS combined,

P=0.15). Furthermore, we do not find any significant enrichment in strong sweeps signals within the nSL or iHS top 1000 (P=0.77), as we do at CoV-VIPs (Figure 1). When considered individually rather than all together, only four of the 20 functions have a significant sweep enrichment (P<0.05; Table S2).

To test whether these four functions explain the sweep enrichment at CoV-VIPs, we test this sweep enrichment at CoV-VIPs again, but this time excluding all genes included in the four previous GO annotations. The sweep enrichment at the remaining CoV-VIPs (91% of them) is the same as when testing all CoV-VIPs (the sum of differences between observed and expected numbers over all the nSL sweep rank thresholds, over all window sizes, and over all five East Asian populations is 14,620 for 352 CoV-VIPs included in the test, versus 15,848 for 385 CoV-VIPs included when not excluding GO functions, in other words almost perfectly proportional to the number of genes included in the test), thus showing that these four host functions do not explain the sweep enrichment at CoV-VIPs. Moreover, further excluding all genes with the 20 GO annotations over-represented more than twofold at CoV-VIPs, we find that the remaining CoV-VIPs (58% of them) have a stronger sweep enrichment than when considering all CoV-VIPs (the sum of differences between observed and expected numbers is 10,843 for 222 genes included in this test, proportionally more than the 15,848 sum of differences for 385 CoV-VIPs when not excluding GO functions). Excluding the genes with any of 106 over-represented GO annotations at CoV-VIPs, we also find that the remaining CoV-VIPs (16% of them) have a stronger sweep enrichment than when considering all CoV-VIPs (sum of differences 4,575 for 62 CoV-VIPs). Host intrinsic functions, as annotated by GO, thus cannot explain the sweep enrichment at CoV-VIPs.

Nevertheless, we further test which GO annotations enriched at CoV-VIPs have a significant peak of Relate times around 900 generations ago, as we did before for CoV-VIPs. To do this, we consider all GO annotations enriched at COV-VIPs, but this time compared to completely random controls, rather than compared to control sets matched for confounding factors as before. Indeed, we previously tested the significance of the peak around 900 generations ago at CoV-VIPs compared to completely random controls (Methods), and here we do the same for a fair comparison. Compared to fully random controls, CoV-VIPs are significantly enriched in 316 GO annotations (P<0.001). Of these 316 GO annotations, 238 are enriched more than two-fold, many more than the 20 GO annotations enriched more than two-fold when using controls matched for confounding factors. This shows that controlling for the confounding factors that we take into account (STAR Methods) effectively controls for many other correlated host intrinsic functions. A total of 39 GO annotations are enriched more than four fold at CoV-VIPs when compared to fully random controls. When considered all together, all the 1,134 genes in the genome other than CoV-VIPs, but with at least one of these 39 highly enriched GO annotations, do not have a significant peak of Relate times (peak significance test P=0.18). When considered individually, only 16 of all the initial 316 over-represented GO annotations have a significant peak between 770 and 970 generations ago (Table S3; peak significance test P<0.05). When removing all CoV-VIPs with at least one of these 16 GO annotations (31% of them), the magnitude of the peak around 900 generations ago at the remaining CoV-VIPs compared to all CoV-VIPs is not affected (Figure S1A,B).

Taken together, these results make it very unlikely that host intrinsic functions explain the patterns and timing of adaptation observed at CoV-VIPs, and make a causal role of coronavirus-like viruses more plausible. Below, we provide further, virus-focused functional evidence, further supporting this.

### Distribution of the interactions of peak-VIPs with different SARS-CoV-2 viral proteins

To further assess the plausibility of an ancient coronavirus-like epidemic, we further analyse the peak-VIPs compared to other CoV-VIPs, but this time by looking at the SARS-CoV-2 viral proteins that they interact with, rather than their functional characteristics in the host. If an ancient epidemic was caused by a coronavirus-like virus with viral proteins orthologous to SARS-CoV-2 proteins, then there is a chance that the distribution of the interactions of the peak-VIPs with the 26 different SARS-CoV-2 proteins included in the Gordon et al. host-virus interactome, may deviate from random expectations, and not simply reflect the number of SARS-CoV-2 VIPs that each SARS-CoV-2 protein interacts with. For example, the peak-VIPs may interact more than expected with specific SARS-CoV-2 proteins, or less than expected. Such deviations from randomness would provide further indirect functional evidence in support of an ancient epidemic, since no specific pattern of interaction with specific viral proteins is expected if adaptation at CoV-VIPs is only a fortuitous coincidence and not caused by an ancient virus.

In total, 35 of the 42 peak-VIPs are SARS-CoV-2 VIPs (Tables S1 and S5). Of the 26 SARS-CoV-2 proteins included in the host-virus interactome identified by Gordon et al., only the NSP12 protein, the most crucial polymerase protein for viral replication, has marginally more interacting peak-VIPs than expected by chance compared to other CoV-VIPs (5 vs. 2.1 expected, permutation test P=0.045; Table S5). A more remarkable, collective pattern is however visible. NSP12 set aside, a high number of 13 SARS-CoV-2 proteins interact with one, and only one peak-VIP (Table S5). This pattern of interaction is more unexpected (13 viral proteins with only one peak-VIP vs. 6.2 expected, permutation test P=0.0026) than the number of peak-VIPs interacting with NSP12, and can be described as an over-dispersion of single peak-VIPs between different SARS-CoV2 viral proteins. This overdispersion is unexpected and a clear departure from randomness, and further strengthens the case for an ancient coronavirus-like epidemic in East Asia.

### Supplemental Tables

**Table S1. VIPs used in the analysis**
Provided as a separate supplemental file. This table provides the Ensembl gene IDs of the VIPs used in the analysis. All VIPs tab: all the 5,291 VIPs annotated to date. RNA VIPs tab: all the 3,952 VIPs annotated to date. CoV-VIPs tab: all the 424 CoV-VIPs annotated to date, with HGNC symbols. SARS-CoV-2 VIPs tab: all the 332 SARS-CoV-2 VIPs annotated to date, with their HGNC symbols. CoV-VIPs effects tab: all CoV-VIPs with published effects. The Pubmed IDs of the papers reporting effects, the corresponding viruses, the altered viral function, and the antiviral or proviral effects are reported in a consistent order in their respective columns, and separated by commas when multiple viruses have antiviral or proviral effects. Peak-VIPs effects tab: Same as CoV-VIPs effects tab, but for peak VIPs only.

| GO term ID | GO term | enrichment curve P, top 10,000 to top 10 nSL ranks, all window sizes | enrichment curve P, top 1,000 to top 10 nSL ranks, 1Mb+2Mb window sizes |
|---|---|---|---|
| GO:0005788 | endoplasmic reticulum lumen | 0.35 | 0.57 |
| GO:0005793 | endoplasmic reticulum-Golgi intermediate compartment | 0.35 | 0.87 |
| GO:0006457 | protein folding | 0.02 | <0.01 |
| GO:0007249 | I-kappaB kinase/NF-kappaB signaling | 0.55 | 0.86 |
| GO:0009100 | glycoprotein metabolic process | 0.08 | 0.35 |
| GO:0009408 | response to heat | 0.55 | 0.8 |
| GO:0010256 | endomembrane system organization | 0.13 | 0.6 |
| GO:0016441 | posttranscriptional gene silencing | 0.15 | 0.78 |
| GO:0016758 | transferase activity, transferring hexosyl groups | <0.01 | 0.33 |
| GO:0016859 | cis-trans isomerase activity | 0.3 | 0.72 |
| GO:0019003 | GDP binding | <0.01 | 0.13 |
| GO:0019058 | viral life cycle | 0.87 | 0.45 |
| GO:0019866 | organelle inner membrane | 0.62 | 0.83 |
| GO:0031047 | gene silencing by RNA | 0.13 | 0.55 |
| GO:0034605 | cellular response to heat | 0.65 | 0.65 |
| GO:0035194 | post-transcriptional gene silencing by RNA | 0.2 | 0.68 |
| GO:0035195 | gene silencing by miRNA | 0.1 | 0.68 |
| GO:0036503 | ERAD pathway | 0.05 | 0.27 |
| GO:0044766 | multi-organism transport | 0.52 | 0.6 |
| GO:1902579 | multi-organism localization | 0.46 | 0.53 |

**Table S2. Gene Ontology terms enriched at CoV-VIPs compared to confounding factors-matched controls.**
Related to Figure 1.

| GO term ID | GO term |
|---|---|
| GO:0000280 | nuclear division |
| GO:0006412 | translation |
| GO:0006414 | translational elongation |
| GO:0006518 | peptide metabolic process |
| GO:0009056 | catabolic process |
| GO:0009266 | response to temperature stimulus |
| GO:0010243 | response to organonitrogen compound |
| GO:0030010 | establishment of cell polarity |
| GO:0043043 | peptide biosynthetic process |
| GO:0043603 | cellular amide metabolic process |
| GO:0044248 | cellular catabolic process |
| GO:0048285 | organelle fission |
| GO:0051235 | maintenance of location |
| GO:1901565 | organonitrogen compound catabolic process |
| GO:1901575 | organic substance catabolic process |
| GO:1901698 | response to nitrogen compound |

**Table S3. Gene Ontology terms with a significant peak between 770 and 970 generations**
Related to Figure 2.

| Ensembl gene ID | HGNC symbol | chromosome, start, end | CDX nSL rank | CHB nSL rank | CHS nSL rank | JPT nSL rank | KHV nSL rank |
|---|---|---|---|---|---|---|---|
| ENSG00000014824 | SLC30A9 | 4 41992489 42089551 | 203 | 213 | 132 | 146 | 86 |
| ENSG00000067560 | RHOA | 3 49396578 49450431 | 24 | 33 | 20 | 120 | 28 |
| ENSG00000082805 | ERC1 | 12 1099675 1605090 | 115 | 120 | 98 | 4 | 106 |
| ENSG00000102898 | NUTF2 | 16 67880635 67906470 | 290 | 155 | 83 | 198 | 206 |
| ENSG00000114302 | PRKAR2A | 3 48782030 48885279 | 15099 | 100 | 61 | 135 | 3538 |
| ENSG00000117054 | ACADM | 1 76190036 76253260 | 55 | 215 | 340 | 667 | 167 |
| ENSG00000126001 | CEP250 | 20 34042985 34099804 | 378 | 87 | 237 | 34 | 671 |
| ENSG00000131043 | AAR2 | 20 34824381 34858840 | 583 | 167 | 148 | 85 | 446 |
| ENSG00000135968 | GCC2 | 2 109065017 109125871 | 3 | 1 | 14 | 5 | 4 |
| ENSG00000154240 | CEP112 | 17 63631656 64188202 | 162 | 430 | 180 | 201 | 145 |
| ENSG00000166794 | PPIB | 15 64448011 64455404 | 5 | 13 | 33 | 123 | 6 |
| ENSG00000169972 | PUSL1 | 1 1243947 1247057 | 63 | 205 | 186 | 582 | 69 |
| ENSG00000178035 | IMPDH2 | 3 49061758 49066841 | 713 | 14 | 4 | 51 | 90 |
| ENSG00000179562 | GCC1 | 7 127220672 127233665 | 14 | 83 | 162 | 22 | 21 |
| ENSG00000204256 | BRD2 | 6 32936437 32949282 | 114 | 900 | 775 | 406 | 925 |
| ENSG00000204435 | CSNK2B | 6 31633013 31638120 | 766 | 375 | 2339 | 348 | 194 |
| ENSG00000204463 | BAG6 | 6 31606805 31620482 | 793 | 338 | 2272 | 366 | 186 |
| ENSG00000233276 | GPX1 | 3 49394609 49396033 | 30 | 43 | 24 | 133 | 31 |
| ENSG00000101346 | POFUT1 | 20 30795683 30826470 | 1405 | 408 | 647 | 4713 | 101 |
| ENSG00000136485 | DCAF7 | 17 61627822 61671639 | 183 | 191 | 182 | 4690 | 377 |
| ENSG00000171552 | BCL2L1 | 20 30252255 30311792 | 10183 | 539 | 2167 | 3203 | 141 |
| ENSG00000204386 | NEU1 | 6 31825436 31830683 | 131 | 236 | 499 | 213 | 314 |
| ENSG00000204536 | CCHCR1 | 6 31110216 31126015 | 847 | 419 | 2502 | 477 | 155 |
| ENSG00000213722 | DDAH2 | 6 31694815 31698394 | 147 | 223 | 497 | 202 | 287 |

**Table S4. CoV-VIPs within the top 200 nSL for at least one of the five East Asian populations**

CDX: Chinese Dai. CHB: Han Chinese from Beijing. CHS: Southern Han Chinese. JPT: Japanese from Tokyo. KHV: Vietnamese Kinh from Hanoi. Related to Figure 1.

| Ensembl gene ID | HGNC symbol | chromosome, gene start, end | Relate selection start | Related selected mutation coordinate | average nSL rank | SARS-CoV-2 VIP | interacts with SARS-CoV-2 |
|---|---|---|---|---|---|---|---|
| ENSG00000023318 | ERP44 | 9 102741461 102861322 | 893.61 | 102806266 | 12026.8 | yes | orf8 |
| ENSG00000047315 | POLR2B | 4 57843888 57897334 | 852.36 | 58572656 | 11682.4 | no | |
| ENSG00000063046 | EIF4B | 12 53399942 53435993 | 833.92 | 52586454 | 8619.2 | no | |
| ENSG00000067560 | RHOA | 3 49396578 49450431 | 770.87 | 50288243 | 45 | yes | nsp7 |
| ENSG00000068912 | ERLEC1 | 2 54014181 54045956 | 847.38 | 53370528 | 12469.4 | yes | orf8 |
| ENSG00000074800 | ENO1 | 1 8921061 8939308 | 778.86 | 8149324 | 3449.8 | no | |
| ENSG00000084733 | RAB10 | 2 26256976 26360323 | 945.68 | 26364468 | 1103.6 | yes | nsp7 |
| ENSG00000102471 | NDFIP2 | 13 80055287 80130210 | 806.41 | 79720040 | 13226.8 | yes | orf9c |
| ENSG00000106344 | RBM28 | 7 127950437 127983962 | 965.75 | 127831580 | 1840.4 | yes | N |
| ENSG00000107929 | LARP4B | 10 855484 977564 | 909.25 | 788658 | 4643.4 | yes | nsp12 |
| ENSG00000114302 | PRKAR2A | 3 48782030 48885279 | 804.56 | 49806393 | 3786.6 | yes | nsp13 |
| ENSG00000114745 | GORASP1 | 3 39138150 39149854 | 936.56 | 39639754 | 8018.6 | yes | nsp13 |
| ENSG00000115310 | RTN4 | 2 55199325 55339757 | 931.35 | 55196956 | 8869.2 | yes | M |
| ENSG00000117054 | ACADM | 1 76190036 76253260 | 862.1 | 76111796 | 288.8 | yes | M |
| ENSG00000117616 | RSRP1 | 1 25568728 25664704 | 913.09 | 26398582 | 15050.2 | no | |
| ENSG00000119782 | FKBP1B | 2 24272571 24286551 | 871.49 | 23766813 | 8879.2 | no | |
| ENSG00000131043 | AAR2 | 20 34824381 34858840 | 863.08 | 35626627 | 285.8 | yes | M |
| ENSG00000131238 | PPT1 | 1 40538379 40563375 | 900.94 | 41272905 | 13419.8 | yes | orf10 |
| ENSG00000134809 | TIMM10 | 11 57295936 57298276 | 840.36 | 57212749 | 1968.2 | yes | nsp4 |
| ENSG00000135968 | GCC2 | 2 109065017 109125871 | 884.89 | 109082052 | 5.4 | yes | nsp13 |
| ENSG00000137073 | UBAP2 | 9 33921691 34048947 | 889.34 | 34030379 | 14631 | yes | nsp12 |
| ENSG00000137409 | MTCH1 | 6 36935917 36954074 | 936.14 | 36107172 | 8763 | yes | orf6 |
| ENSG00000138698 | RAP1GDS1 | 4 99182535 99365012 | 770.33 | 100056998 | 4412.4 | yes | nsp2 |
| ENSG00000138829 | FBN2 | 5 127593601 127994878 | 967.68 | 127786577 | 1337.8 | yes | nsp9 |
| ENSG00000140577 | CRTC3 | 15 91073157 91188577 | 924.95 | 91311017 | 14700.4 | yes | nsp12 |
| ENSG00000143653 | SCCPDH | 1 246887349 246931439 | 893.04 | 246816033 | 3280.4 | yes | nsp7 |
| ENSG00000156599 | ZDHHC5 | 11 57435219 57468659 | 840.36 | 57212749 | 1440.4 | yes | Spike |
| ENSG00000158545 | ZC3H18 | 16 88636789 88698374 | 923.5 | 88801026 | 5751.2 | yes | E |
| ENSG00000164609 | SLU7 | 5 159828648 159848718 | 910.65 | 159975072 | 10859.4 | yes | nsp12 |
| ENSG00000165527 | ARF6 | 14 50359810 50361490 | 958.32 | 50389457 | 1255.8 | yes | nsp15 |
| ENSG00000165661 | QSOX2 | 9 139098179 139137687 | 863.7 | 139236401 | 5229.2 | yes | nsp7 |
| ENSG00000165688 | PMPCA | 9 139305110 139318213 | 863.7 | 139236401 | 3526 | yes | M |
| ENSG00000166794 | PPIB | 15 64448011 64455404 | 784.21 | 64924315 | 36 | no | |
| ENSG00000166949 | SMAD3 | 15 67356101 67487533 | 866.1 | 68202469 | 7934.4 | no | |
| ENSG00000167461 | RAB8A | 19 16222439 16245044 | 818.26 | 17037080 | 13110.6 | yes | nsp7 |
| ENSG00000169972 | PUSL1 | 1 1243947 1247057 | 854.29 | 1226292 | 221 | yes | orf8 |
| ENSG00000177917 | ARL6IP6 | 2 153574407 153617688 | 782.85 | 154026994 | 4375.2 | yes | orf3a |
| ENSG00000178035 | IMPDH2 | 3 49061758 49066841 | 804.56 | 49806393 | 174.4 | yes | nsp14 |
| ENSG00000179562 | GCC1 | 7 127220672 127233665 | 965.75 | 127831580 | 60.4 | yes | nsp13 |
| ENSG00000230989 | HSBP1 | 16 83841448 83853342 | 878.92 | 84827154 | 2213.2 | yes | nsp13 |
| ENSG00000233276 | GPX1 | 3 49394609 49396033 | 770.87 | 50288243 | 52.2 | yes | nsp5_C145A |
| ENSG00000240344 | PPIL3 | 2 201735630 201754026 | 866.27 | 201527338 | 11401.4 | yes | nsp12 |

**Table S5. The 42 peak-VIPs with selection start times between 770 and 970 generations** Relate selection start times are the average selection start time over the five East Asisan populations. The average nSL ranks are the average over the five East Asian populations, for 1Mb windows (STAR Methods). Related to Figure 2.

| HGNC symbol | SNP ID | Africa | Europe | East Asia |
|---|---|---|---|---|
| ERP44 | rs1418268 | 0.006 | 0.084 | 0.496 |
| POLR2B | rs34276339 | 0.041 | 0.302 | 0.517 |
| EIF4B | rs4762061 | 0.006 | 0.062 | 0.505 |
| RHOA | rs201632611 | 0.001 | 0.012 | 0.556 |
| ERLEC1 | rs1374254 | 0.013 | 0.15 | 0.57 |
| ENO1 | rs11121084 | 0.047 | 0.003 | 0.541 |
| RAB10 | rs76245846 | 0.002 | 0.004 | 0.665 |
| NDFIP2 | rs9565467 | 0 | 0.05 | 0.45 |
| RBM28 | rs896183 | 0.041 | 0.408 | 0.624 |
| LARP4B | rs72635991 | 0.112 | 0.245 | 0.547 |
| PRKAR2A | rs57704135 | 0.001 | 0.006 | 0.617 |
| GORASP1 | rs11129839 | 0.016 | 0.108 | 0.589 |
| RTN4 | rs2580764 | 0.039 | 0.398 | 0.621 |
| ACADM | rs554189868 | 0.032 | 0.209 | 0.626 |
| RSRP1 | rs113496569 | 0.018 | 0.219 | 0.513 |
| FKBP1B | rs1877789 | 0.003 | 0.013 | 0.64 |
| AAR2 | rs6124524 | 0.015 | 0.148 | 0.423 |
| PPT1 | rs76503470 | 0.001 | 0.008 | 0.489 |
| TIMM10 | rs4939145 | 0.199 | 0.407 | 0.552 |
| GCC2 | rs11123695 | 0.002 | 0.011 | 0.851 |
| UBAP2 | rs55660804 | 0.008 | 0.148 | 0.485 |
| MTCH1 | rs2071863 | 0.121 | 0.159 | 0.575 |
| RAP1GDS1 | rs3805322 | 0 | 0 | 0.43 |
| FBN2 | rs72663359 | 0.004 | 0.042 | 0.571 |
| CRTC3 | rs2518967 | 0.023 | 0.239 | 0.539 |
| SCCPDH | rs13374825 | 0.197 | 0.353 | 0.568 |
| ZDHHC5 | rs4939145 | 0.199 | 0.407 | 0.552 |
| ZC3H18 | rs3214056 | 0.155 | 0.181 | 0.554 |
| SLU7 | rs56111320 | 0.095 | 0.053 | 0.493 |
| ARF6 | rs10162475 | 0.192 | 0.136 | 0.558 |
| QSOX2 | rs74312704 | 0.001 | 0.061 | 0.574 |
| PMPCA | rs74312704 | 0.001 | 0.061 | 0.574 |
| PPIB | rs4777495 | 0.005 | 0.025 | 0.62 |
| SMAD3 | rs58511468 | 0.017 | 0.232 | 0.549 |
| RAB8A | rs75106035 | 0.001 | 0.038 | 0.443 |
| PUSL1 | rs2273276 | 0.019 | 0.005 | 0.539 |
| ARL6IP6 | rs2033849 | 0.005 | 0.025 | 0.469 |
| IMPDH2 | rs57704135 | 0.001 | 0.006 | 0.617 |
| GCC1 | rs896183 | 0.041 | 0.408 | 0.624 |
| HSBP1 | rs72628285 | 0.019 | 0.19 | 0.587 |
| GPX1 | rs201632611 | 0.001 | 0.012 | 0.641 |
| PPIL3 | rs2241079 | 0.004 | 0.056 | 0.475 |

**Table S6. Frequencies of the selected alleles in Africa, Europe and East Asia.** Related to Figures 3 and 4.

| **Drugs** | **DrugBank Accession** | **Selected Target** | **COVID-19 Trials** | **Number of Trials** |
|---|---|---|---|---|
| Dexamethasone; Genistein; Leuprolide Acetate; Halofuginone | DB01234 | *SMAD3* | Yes | 10 |
| Mycophenolate Mofetil; Ribavirin; Merimepodib; NAHD | DB00688; DB00811; DB04862 | *IMPDH2* | Yes | 2 |
| Cyclosporine; L-Proline | DB00091 | *PPIB* | Yes | 2 |
| Glutathione | DB00608 | *GPX1* | Yes | 22 |
| Flavin adenine dinucleotide; Octanoyl-Coenzyme A | DB03147; DB02910 | *ACADM* | No | – |
| Myristic Acid; 5'-Guanosine-Diphosphate-Monothiophosphate; Guanosine-5'-Diphosphate | | *ARF6* | No | – |
| Palmitic Acid; Lamotrigin; 1-Hexadecylsulfonyl Fluoride | | *PPT1* | No | – |
| Guanosine-5'-Diphosphate | | *RHOA* | No | – |
| Ozanezumab | | *RTN4* | No | – |
| Druggable Tie3A | (Finan et al., 2017) | *ERP44* | No | – |
| Druggable Tie3A | (Finan et al., 2017) | *FBN2* | No | – |
| Druggable Tie1 | (Finan et al., 2017) | *FKBP1B* | No | – |
| Druggable Tie1 | (Finan et al., 2017) | *PPIL3* | No | – |
| Druggable Tie3A | (Finan et al., 2017) | *QSOX2* | No | – |
| Druggable Tie3B | (Finan et al., 2017) | *AAR2* | No | – |

**Table S7. Drugs targeting the 42 peak-VIPs.**
Related to Figure 2.

### Supplemental Figures

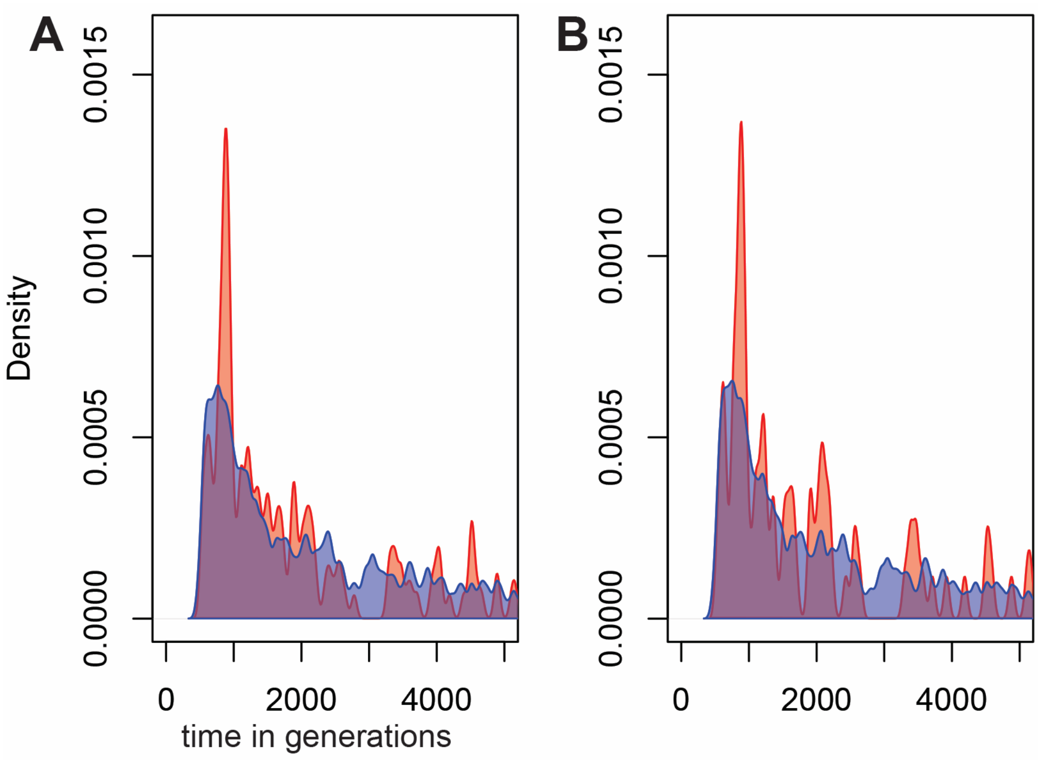


**Figure S1. Timing of selection start at CoV-VIPs, with or without removing GO functions with a significant peak between 770 and 970 generations**
Same legend as Figure 2. A) All CoV-VIPs. B) CoV-VIPs with at least one of the 16 GO functions with a significant peak between 770 and 970 generations ago are excluded (31% of VIPs; Table S3). Related to Figure 2.

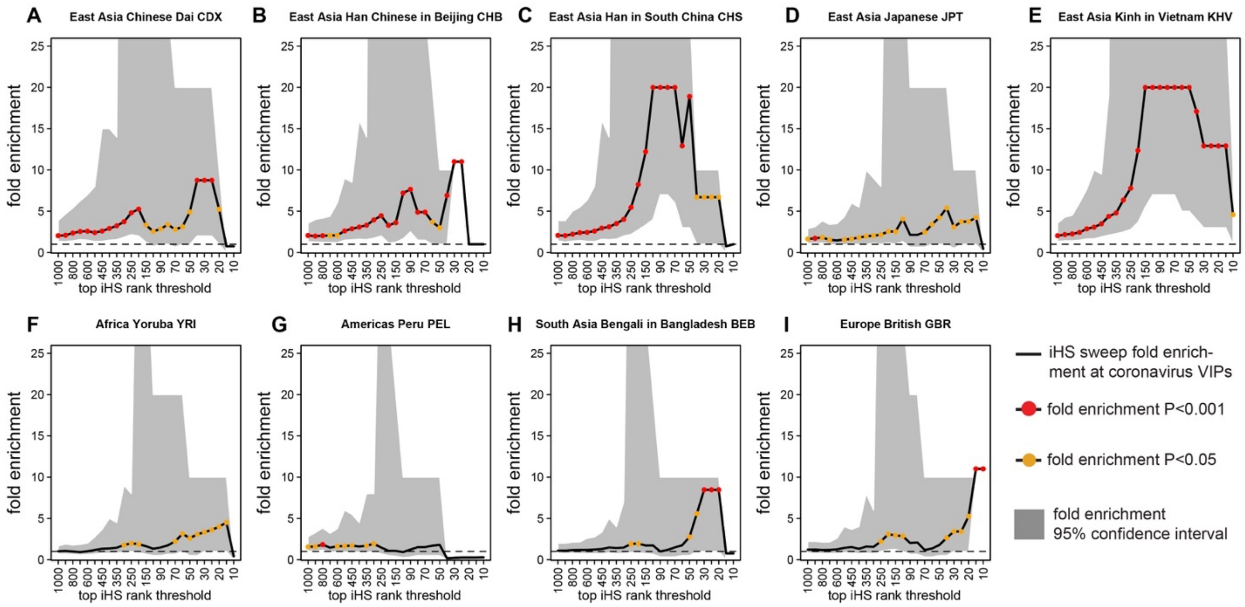


**Figure S2. CoV-VIPs sweep enrichment with iHS**

Same legend as Figure 1. The only change compared to Figure 1 is the use of his instead of nSL. Related to Figure 1.

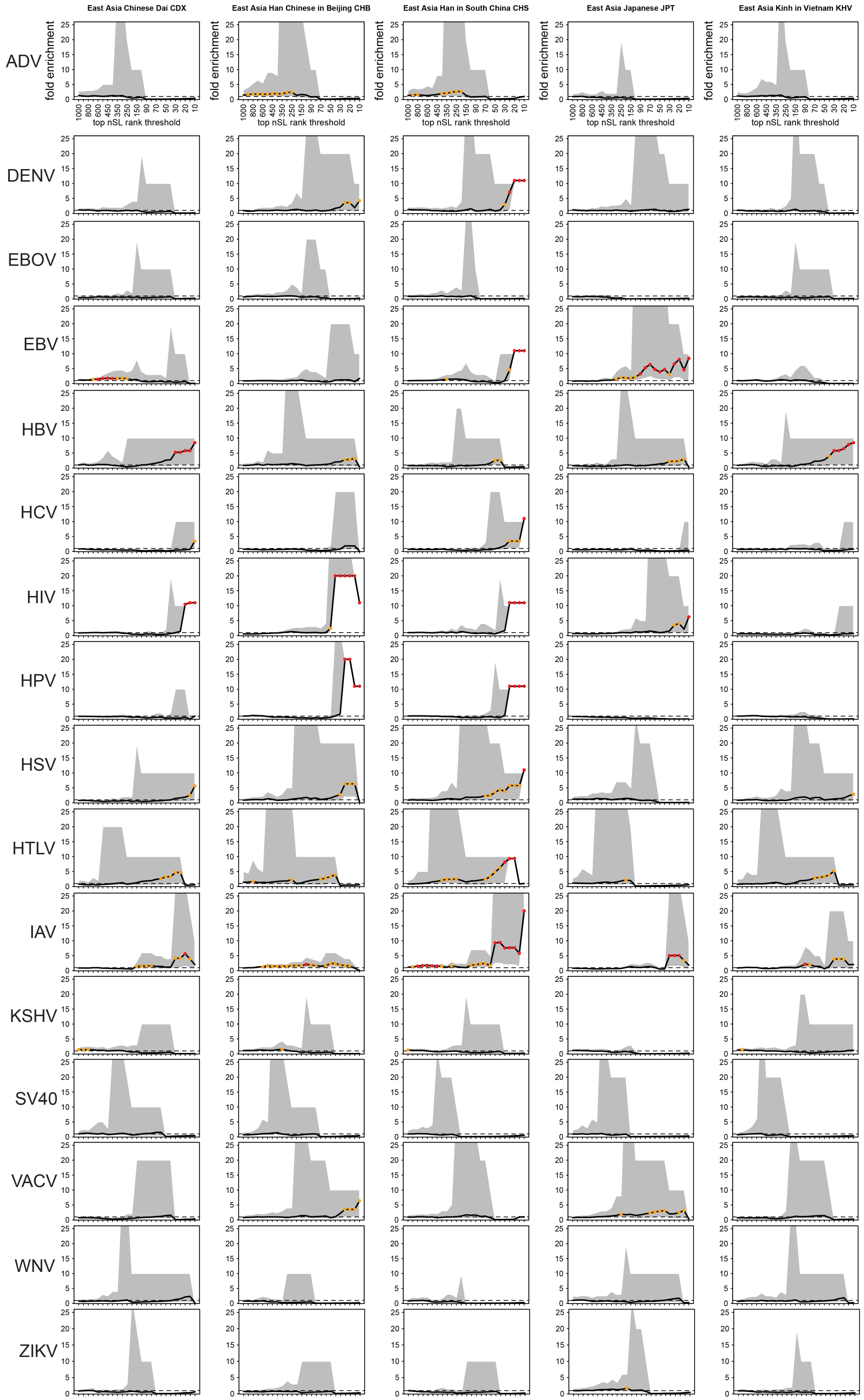


**Figure S3. nSL sweep enrichment curves for 17 other viruses in East Asia**
Same legend as in Figure 1. Whole curve P>0.05 for all viruses. Related to Figure 1.

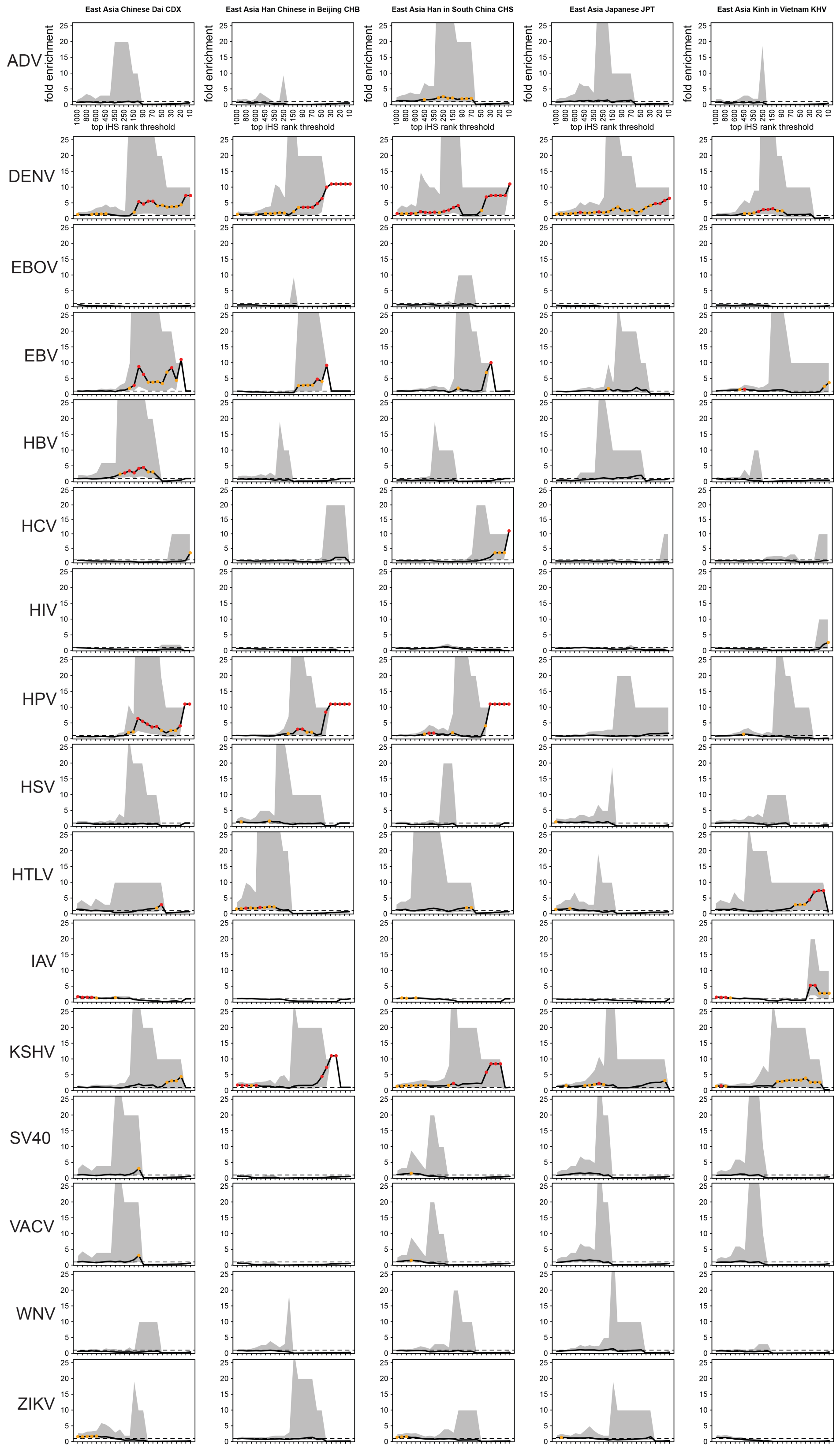


**Figure S4. iHS sweep enrichment curves for 17 other viruses in East Asia**

Same legend as in Figure 1. Whole curve P>0.05 for all viruses. Related to Figure 1.

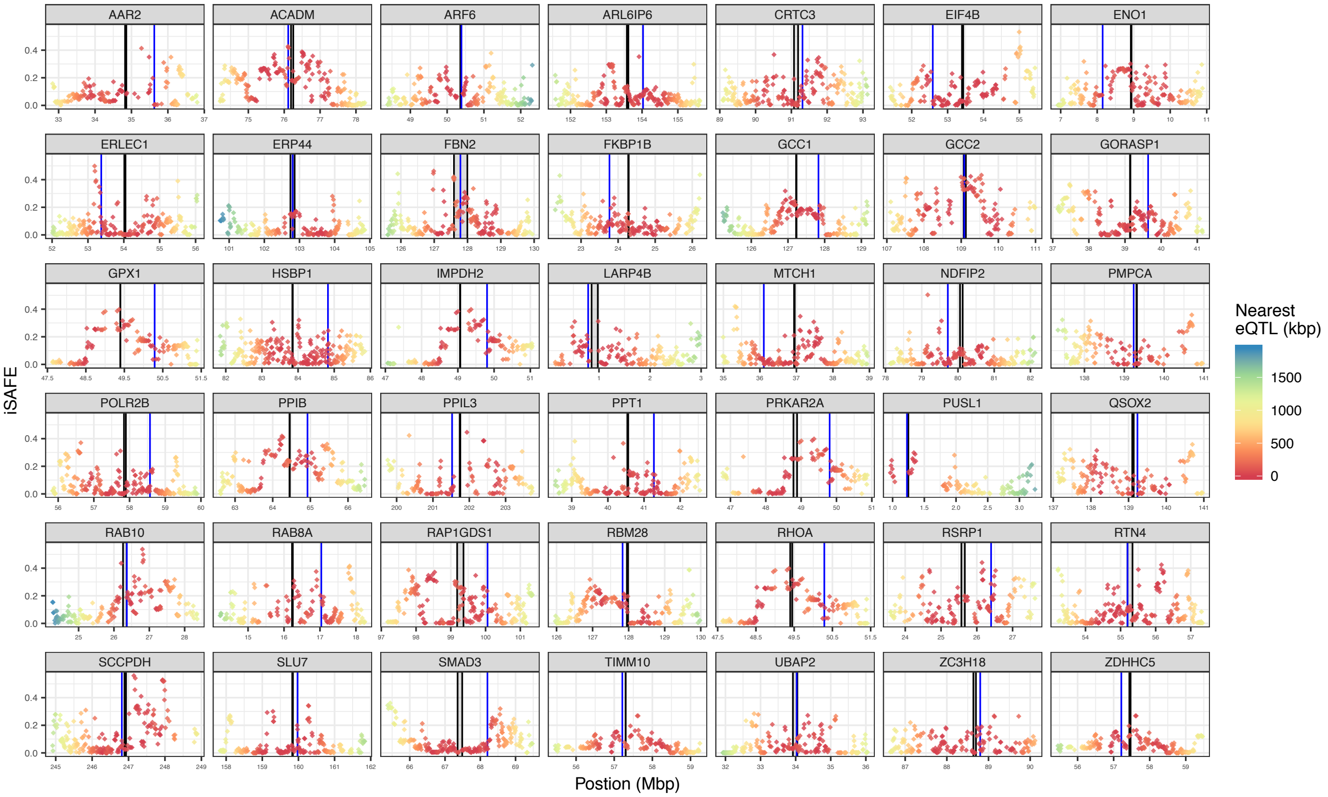


**Figure S5. iSAFE peaks, GTEx eQTLs and Relate selected variants locations**
Dark lines: gene starts and gene ends. Blue line: Relate selected variant location. The color scale provides information about distance to the nearest GTEx eQTL. iSAFE peaks are not always clean, sharp peaks, and the Relate selected variants do not always overlap local iSAFE peaks, possibly as a result of both recombination since strong selection stopped, and weaker selection in more recent times (Figure 4). This is suggested by multiple steep iSAFE drops in the middle of peaks, as visible for example for ARL6IP6 at coordinate 153Mb. Related to Figure 5.

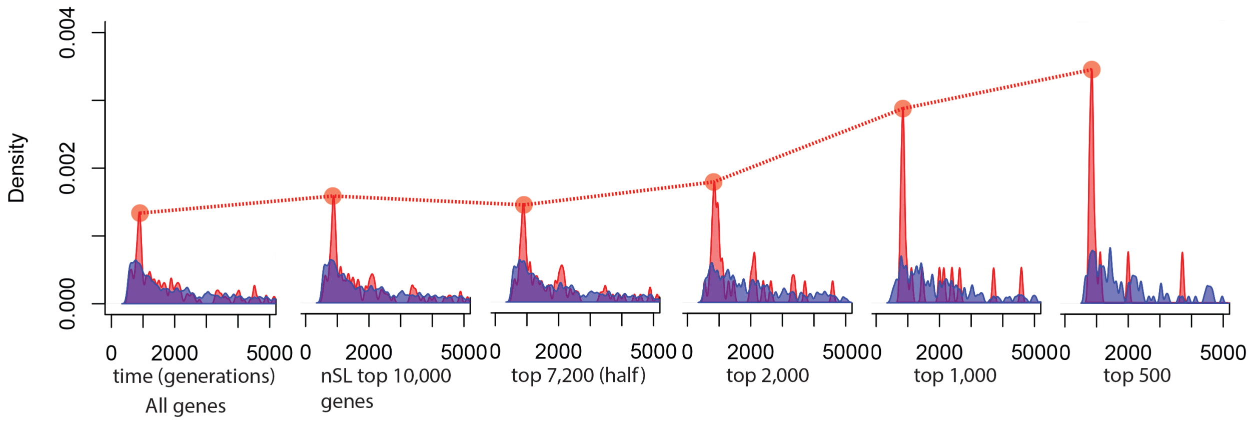


**Figure S6. Selection timing peak for increasingly stronger nSL sets of genes**
Legend as for Figure 2. The figure shows the amplitude of the peak of selection start times for increasingly high nSL thresholds. For example, for the nSL top 1,000, only selection start times at genes within the top 1,000 nSL (average rank over East Asian populations, lower rank of the 1Mb and 2Mb nSL windows) are included to get the pink and blue distributions. Related to Figure 2.

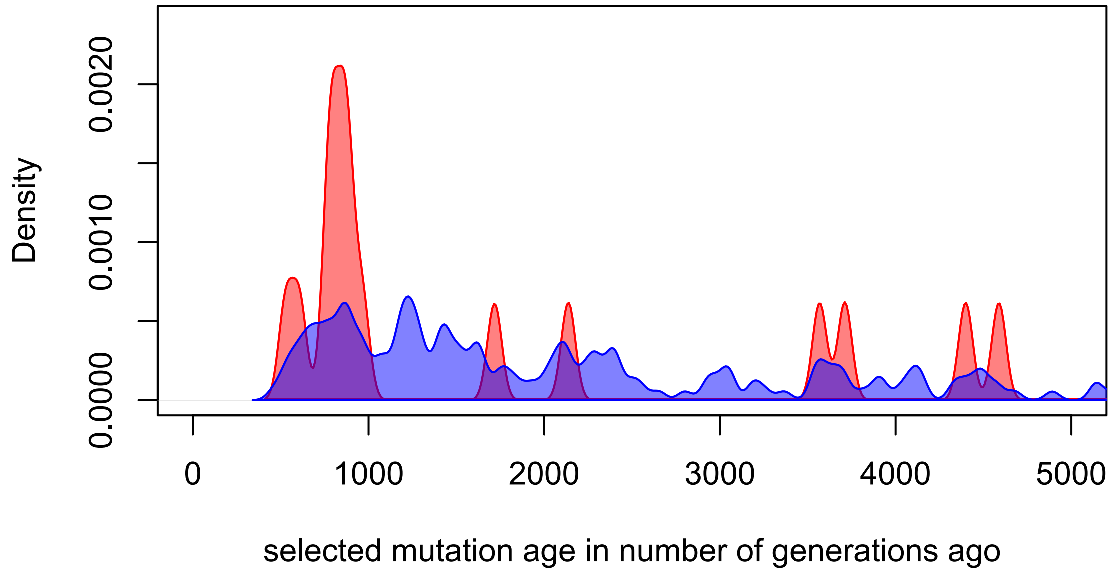


**Figure S7. Selection timing at top Tajima's D genes**
Legend as for Figure 2. The two distributions include only genes within Tajima's D top 1,000 (rank averaged over the five East Asian populations, Tajima's D measured in 1Mb windows). Related to Figure 2.

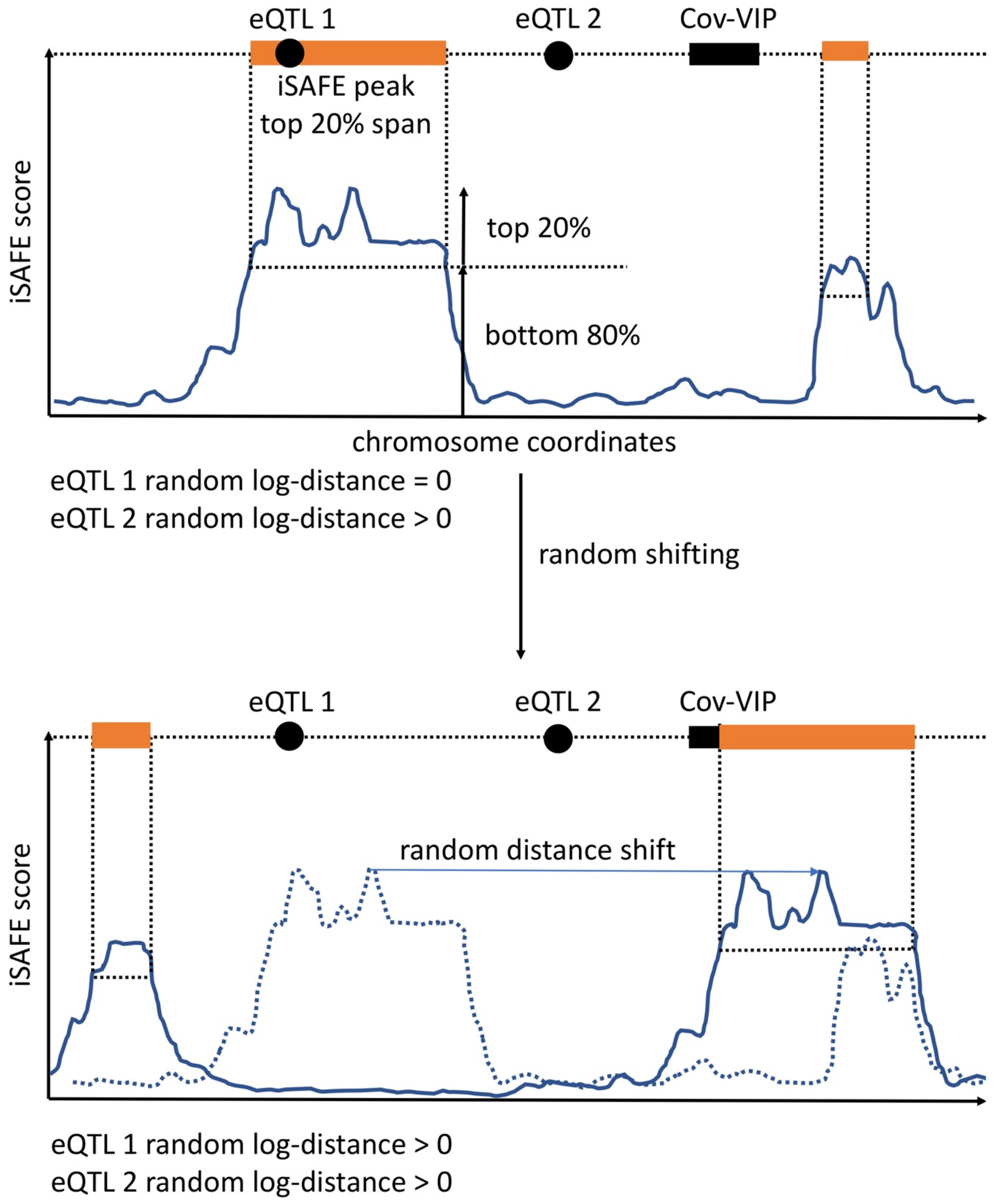


**Figure S8. Sliding of iSAFE coordinates for the proximity ratio test**
Black rectangle: area between the transcription start and end of a CoV-VIP. Black dot: coordinate of eQTL for the corresponding CoV-VIP. Orange area: area where distance between the iSAFE peak area and the closest eQTL is counted as zero. If the eQTL falls outside of an orange area, the distance is counted as distance to closest orange area edge. Dashed blue line in thre lower panel: original location of the real iSAFE score before random sliding. Related to Figure 5.
